# Adolescent social anxiety is associated with diminished discrimination of anticipated threat and safety in the bed nucleus of the stria terminalis

**DOI:** 10.1101/2023.10.30.564701

**Authors:** Juyoen Hur, Rachael M. Tillman, Hyung Cho Kim, Paige Didier, Allegra S. Anderson, Samiha Islam, Melissa D. Stockbridge, Andres De Los Reyes, Kathryn A. DeYoung, Jason F. Smith, Alexander J. Shackman

**Affiliations:** Department of Psychology, Yonsei University, Seoul 03722, Republic of Korea; Department of Neuropsychology, Children’s National Hospital, Washington, DC 20010 USA; Department of Psychology, University of Maryland, College Park, MD 20742 USA; Neuroscience and Cognitive Science Program, University of Maryland, College Park, MD 20742 USA; Maryland Neuroimaging Center, University of Maryland, College Park, MD 20742 USA; TheraQuest LLC, Bethesda, MD 20817; Department of Psychological Sciences, Vanderbilt University, Nashville, TN 37240 USA; Department of Psychology, University of Pennsylvania, Philadelphia, PA 19104 USA; Department of Neurology, School of Medicine, Johns Hopkins University, Baltimore, MD 21287 USA

**Keywords:** bed nucleus of the stria terminalis (BST/BNST), central extended amygdala, developmental affective neuroscience, developmental psychopathology, pediatric social anxiety, social phobia

## Abstract

Social anxiety—which typically emerges in adolescence—lies on a continuum and, when extreme, can be devastating. Socially anxious individuals are prone to heightened fear, anxiety, and the avoidance of contexts associated with potential social scrutiny. Yet most neuroimaging research has focused on acute social threat. Much less attention has been devoted to understanding the neural systems recruited during the uncertain anticipation of potential encounters with social threat. Here we used a novel fMRI paradigm to probe the neural circuitry engaged during the anticipation and acute presentation of threatening faces and voices in a racially diverse sample of 66 adolescents selectively recruited to encompass a range of social anxiety and enriched for clinically significant levels of distress and impairment. Results demonstrated that adolescents with more severe social anxiety symptoms experience heightened distress when anticipating encounters with social threat, and reduced discrimination of uncertain social threat and safety in the bed nucleus of the stria terminalis (BST), a key division of the central extended amygdala (EAc). Although the EAc—including the BST and central nucleus of the amygdala—was robustly engaged by the acute presentation of threatening faces and voices, the degree of EAc engagement was unrelated to the severity of social anxiety. Together, these observations provide a neurobiologically grounded framework for conceptualizing adolescent social anxiety and set the stage for the kinds of prospective-longitudinal and mechanistic research that will be necessary to determine causation and, ultimately, to develop improved interventions for this often-debilitating illness.

## INTRODUCTION

Socially anxious individuals are prone to heightened fear, anxiety, and the avoidance of contexts associated with the potential for social scrutiny (<u>Beidel et al., 2019</u>). Social anxiety lies on a continuum and, when extreme, can be devastating, with functional impairment evident in many individuals who do not meet full diagnostic criteria (<u>Fehm et al., 2008</u>; <u>Hyett & McEvoy, 2018</u>; <u>Katzelnick et al., 2001</u>; <u>Kessler, 2003</u>; <u>Lipsitz & Schneier, 2000</u>; <u>Merikangas et al., 2002</u>; <u>Schneier et al., 2002</u>). Social anxiety disorder (SAD) is among the most common psychiatric illnesses (lifetime prevalence: ∼13%), typically emerges in adolescence, and confers heightened risk for a variety of other developmental, academic, and psychiatric problems, including co-morbid internalizing disorders and substance misuse (<u>Angst et al., 2016</u>; <u>Beesdo-Baum & Knappe, 2012</u>; <u>Ernst et al., 2023</u>; <u>Gregory et al., 2007</u>; <u>Hyett & McEvoy, 2018</u>; <u>Jystad et al., 2021</u>; <u>Kessler et al., 2012</u>; <u>Koyuncu et al., 2019</u>; <u>Mathew et al., 2011</u>; <u>Schneier et al., 1992</u>; <u>Stein et al., 2017</u>). Existing treatments are inconsistently effective or associated with significant adverse effects, underscoring the urgency of developing a more complete understanding of the neural systems governing social anxiety in the first decades of life (<u>Batelaan et al., 2017</u>; <u>Beidel et al., 2019</u>; <u>Cuijpers et al., 2024</u>; <u>Evans et al., 2021</u>; <u>James et al., 2020</u>; <u>Rapee et al., 2023</u>; <u>Scholten et al., 2016</u>; <u>Scholten et al., 2013</u>; <u>Singewald et al., 2023</u>; <u>Spinhoven et al., 2016</u>; <u>Strawn et al., 2021</u>).

Socially anxious adults and youth are prone to elevated distress and arousal in two distinct contexts: **(a)** when social threat is acute (i.e., certain and imminent), as when performing in front of a group; and **(b)** when social threat is possible, but uncertain in timing or likelihood, as when first entering a classroom, conversation, or other social environment (<u>APA, 2022</u>; <u>Cain, 2023</u>; <u>Davis et al., 2010</u>; <u>NIMH, 2011</u>). While the underlying neurobiology is undoubtedly complex and multifactorial, converging lines of preclinical research motivate the hypothesis that socially anxious individuals’ heighted reactivity to both kinds of threat reflect functional alterations in the central extended amygdala (EAc), a neuroanatomical circuit encompassing the dorsal amygdala in the region of the central nucleus (Ce) and the neighboring bed nucleus of the stria terminalis (BST) (<u>Fox, Oler, Tromp, et al., 2015</u>; <u>Fox & Shackman, 2019</u>; <u>Hur et al., 2020</u>; <u>Hur et al., 2019</u>; <u>Moscarello & Penzo, 2022</u>; <u>Shackman & Fox, 2021</u>; <u>Shackman et al., *in press*</u>; <u>Tseng et al., 2023</u>). Both regions are poised to trigger behavioral, psychophysiological, and neuroendocrine responses to threat via dense projections to downstream effector regions (<u>Davis & Whalen, 2001</u>; <u>Fox, Oler, Tromp, et al., 2015</u>). Large-scale neuroimaging studies in monkeys (*n*=238-592) show that Ce and BST metabolism co-varies with trait-like individual differences in freezing, cortisol, and other defensive responses elicited by uncertain naturalistic threats (<u>Fox, Oler, Shackman, et al., 2015</u>; <u>Shackman et al., 2013</u>). Work in humans demonstrates that both regions are sensitive to a broad spectrum of threatening and aversive stimuli (<u>Fox & Shackman, 2019</u>; <u>Hur et al., 2020</u>; <u>Meyer et al., 2019</u>; <u>Murty et al., 2023</u>; <u>Shackman & Fox, 2021</u>). Lesion and other kinds of focal perturbation studies in rodents demonstrate that microcircuits within and between the Ce and BST are critical for orchestrating defensive responses to both acute and uncertain threats (<u>Chen et al., 2022</u>; <u>Fox & Shackman, 2019</u>; <u>Lange et al., 2017</u>; <u>Moscarello & Penzo, 2022</u>; <u>Pomrenze, Giovanetti, et al., 2019</u>; <u>Pomrenze, Tovar-Diaz, et al., 2019</u>; <u>Ren et al., 2022</u>; <u>Ressler et al., 2020</u>; <u>Zhu et al., 2024</u>), and help calibrate the degree of wariness displayed during interactions with unfamiliar and potentially threatening conspecifics (<u>Lee et al., 2008</u>; <u>Lungwitz et al., 2012</u>; <u>Sajdyk et al., 2008</u>). Although the causal contribution of the BST has yet to be explored in primates, monkeys with fiber-sparing lesions of the Ce show reduced defensive responses to both acute (e.g., snake) and uncertain threat (e.g., human intruder’s profile) (<u>Kalin et al., 2004</u>; <u>Oler et al., 2016</u>). Complete destruction of the amygdala is associated with reduced signs of anxiety during interactions with unfamiliar animals (<u>Emery et al., 2001</u>). Likewise, humans with circumscribed amygdala damage show atypically low levels of dispositional fear and anxiety—whether indexed by self-report, family report, clinician report, or daily diary—and display a profound lack of fear and anxiety in response to both acute threats (e.g., spiders, snakes, Pavlovian threat cues, horror movies) and contexts associated with uncertain potential threat (e.g., traversing a haunted house) (<u>Bechara et al., 1995</u>; <u>Feinstein et al., 2011</u>; <u>Feinstein et al., 2016</u>; <u>Korn et al., 2017</u>; <u>Tranel et al., 2006</u>). Conversely, stimulation of the Ce is associated with potentiated responses to uncertain threat in animals and heightened feelings of fear and anxiety in humans, suggesting that circuits centered on this region are necessary and sufficient for many of the core signs and symptoms of social anxiety (<u>Inman et al., 2020</u>; <u>Kalin et al., 2016</u>; <u>Moscarello & Penzo, 2022</u>).

Despite this progress, the relevance of these discoveries to pediatric social anxiety remains unclear. To date, the vast majority of human neuroimaging studies have focused on laboratory analogues of acute social threat, including virtual social interactions and ‘threat-related’ social cues, such as photographs of fearful and angry facial expressions (<u>Jarcho et al., 2013</u>). Meta-analyses show that the EAc is generally hyper-reactive to social cues among adults with SAD and those with a childhood history of extreme shyness and behavioral inhibition, key dispositional risk factors for SAD (<u>Brühl et al., 2014</u>; <u>Chavanne & Robinson, 2021</u>; <u>Clauss & Blackford, 2012</u>; <u>Fox & Kalin, 2014</u>; <u>Gentili et al., 2016</u>; <u>Tan et al., *in press*</u>). Among adults, EAc reactivity to acute social threat is dampened by clinically effective treatments for SAD, consistent with a causal role (<u>Klumpp & Fitzgerald, 2018</u>; <u>Shackman et al., 2016</u>). While the pediatric social anxiety literature is comparatively sparse, the available evidence suggests a broadly consistent pattern (<u>Jarcho et al., 2013</u>). Nevertheless, some comparatively large-scale studies have failed to detect exaggerated EAc reactivity to acute social threat. For example, Ziv and colleagues reported negligible effects of Generalized SAD on amygdala reactivity to dynamically looming hostile faces or film clips of actors delivering social criticism (<u>n=67 unmedicated patients, n=28 matched controls; Ziv et al., 2013</u>).

Much less scientific attention has been devoted to understanding the neural systems underlying heightened reactivity to anticipated social threat. Early work using comparatively low-resolution neuroimaging techniques (e.g., ^15^O-positron emission tomography) and small case-control designs (*n*<10/group) revealed evidence of heightened amygdala activity among adults with SAD during the certain anticipation of a speaking challenge (<u>Lorberbaum et al., 2004</u>; <u>Tillfors et al., 2002</u>)—an effect later confirmed in higher resolution studies (3T fMRI) and larger samples (<u>Boehme et al., 2014</u>; <u>Davies et al., 2017</u>). Potential group differences in BST activation were neither scrutinized nor reported. More recent work has extended this approach to the uncertain anticipation of social threat (<u>Figel et al., 2019</u>). Here a cue presented at the outset of each trial signaled whether the participant’s face would (‘threat’) or would not (‘safety’) be video-recorded. Cues were followed by a temporally variable and uncertain anticipation phase (3-16 s) that terminated with the presentation of a second cue indicating whether the camera was recording. Results revealed sustained levels of heightened BST activation in adults with SAD *and* controls during the uncertain anticipation of social threat encounters. Although patients showed numerically smaller responses, significant group differences were not evident in either the BST or the dorsal amygdala (Ce), and potential regional differences (BST vs. Ce) were not interrogated. While important, the relevance of this body of observations to maladaptive social anxiety earlier in life, during adolescence, remains little explored and largely unknown (<u>but see Michalska et al., 2023 for related efforts in a mixed-anxiety sample of youth</u>).

Here we combined fMRI with a novel threat paradigm—the Maryland Social Threat Countdown (MSTC)— in a racially diverse group of adolescents. The sample was selectively recruited to encompass a broad spectrum of social anxiety—without gaps or discontinuities—and enriched for clinically significant levels of distress and impairment. Adolescent with comorbid anxiety and depressive disorders were enrolled, maximizing clinical relevance (<u>Ernst et al., 2023</u>; <u>Jystad et al., 2021</u>; <u>Koyuncu et al., 2019</u>; <u>Lahey et al., 2022</u>; <u>Tiego et al., 2023</u>). The MSTC task takes the form of a 2 (*Valence:* Social Threat/Safety) × 2 (*Temporal Certainty:* Uncertain/Certain) × 2 (*Phase:* Anticipation/Presentation) randomized event-related design, allowing us to separately assess EAc activation during the anticipation and acute presentation of both certain and uncertain social-threat cues (**Figure 1**). On Certain Social Threat trials, participants saw a descending stream of integers (‘count-down’). To maximize distress, the anticipation period always culminated with the presentation of a multi-modal social threat that included a photograph of an angry young-adult face and an audio clip of a hostile verbal remark (e.g., *“No one wants you here”*). Uncertain Social Threat trials were similar, but the integer stream was randomized and presented for an unsignaled and variable duration. Here, participants knew that a social threat was coming, but they had no way of knowing precisely when the encounter would occur. Safety trials were similar but terminated with the presentation of normatively benign photographs (happy faces) and audio clips (e.g., *“I play soccer”*).

**Figure 1.**
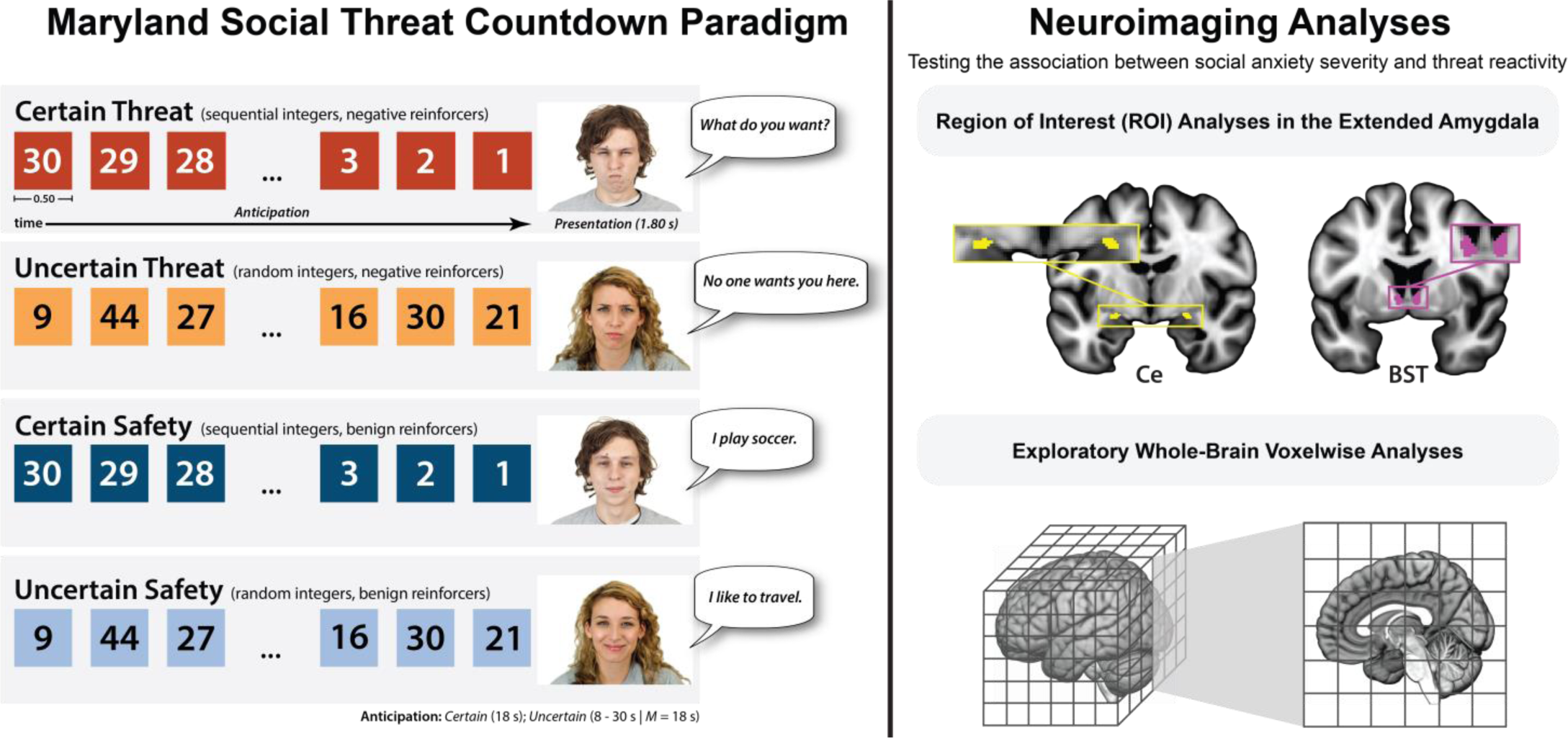
*Study overview.* The Maryland Social Threat Countdown (MSTC) paradigm. As shown schematically in the left panel, the MSTC paradigm takes the form of a 2 (*Valence:* Social Threat/Safety) × 2 (*Temporal Certainty:* Uncertain/Certain) × 2 (*Phase:* Anticipation/Presentation) repeated-measures randomized event-related design. On Certain Social Threat trials, participants saw a descending stream of integers (‘count-down’) for 18 s. To maximize distress, the anticipation epoch always culminated with the presentation of a multi-modal social threat (1.8 s), encompassing a photograph of an angry face and an audio clip of a hostile verbal remark (e.g., *“No one wants you here”*). Uncertain Social Threat trials were similar, but the integer stream was randomized and presented for an uncertain and variable duration (8-30 s; *M*=18 s). Participants knew that a social threat would occur, but they had no way of knowing precisely when the encounter would occur. Safety trials were similar but terminated with the delivery of normatively benign photographs (happy faces) and audio clips (e.g., *“I play soccer”*)**. Neuroimaging analyses.** As shown schematically in the top-right panel, two analytic approaches were used to determine the impact of social anxiety on neural reactivity to social threat. Hypothesis testing focused on two well-established, anatomically defined EAc regions-of-interest (ROIs): the Ce (*yellow*) and BST (*magenta*). Because the ROIs (i.e., voxel-level measurements) were chosen *a priori*—on the basis of neuroanatomy, rather than suprathreshold activation—this approach provides statistically unbiased effect-size estimates (<u>Poldrack et al., 2017</u>). Standardized regression coefficients were extracted and averaged across voxels for each combination of ROI, condition, and participant. Hypothesis testing used a standard general linear model (GLM). For illustrative purposes, 1-mm ROIs are shown. Analyses employed ROIs decimated to the 2-mm resolution of the fMRI data. As shown schematically in the bottom-right panel, exploratory whole-brain voxelwise analyses were also performed. Abbreviations—BST, bed nucleus of the stria terminalis; Ce, central nucleus of the amygdala; *M*, mean; s, seconds.

Anatomically defined regions-of-interest (ROIs) and spatially unsmoothed fMRI data allowed us to rigorously quantify Ce and BST reactivity to the anticipation and presentation of uncertain and certain social threat, and determine whether activation in one or both regions covary with the severity of social anxiety (**Figure 1**). Unlike voxelwise analyses—which screen thousands of voxels for evidence of statistical significance and yield optimistic effect size estimates in suprathreshold regions—anatomically defined ROIs ‘fix’ the outcomes of interest *a priori*, providing statistically unbiased estimates of brain-phenotype associations (<u>Poldrack et al., 2017</u>). Exploratory whole-brain voxelwise analyses were also performed, enabling more direct comparison with prior work.

Discovering the neural systems most relevant to adolescent social anxiety is important. It would provide an empirical rationale for prioritizing specific experimental assays (e.g., the uncertain anticipation of social threat) and neurobiological targets (e.g., BST) for prospective-longitudinal and intervention studies in humans (e.g., acute pharmacological challenges, neurofeedback) and mechanistic studies in animals. Animal models are critical for identifying the molecules and micro-circuits that underlie aberrant EAc threat processing and, ultimately, for developing novel pharmacological treatments for this often-debilitating disorder (<u>Fox & Shackman, 2019</u>).

## METHOD

### Recruitment and Study Overview

Adolescents (13-17 years) were recruited using a multipronged strategy, including online advertisements (e.g., Facebook, parent listservs), flyers posted at community mental health clinics and other high-traffic community settings (e.g., coffee shops, libraries, community centers), and referrals from on-going research studies. To capture a broad spectrum of social anxiety, with substantial enrichment for elevated levels of distress and impairment, separate advertisements were used to target adolescents with elevated (e.g., shy teens) and low-to-middling symptoms. Following preliminary screening, potentially eligible individuals completed a single face-to-face session that included a self-report assessment, a structured clinical interview, and an MRI assessment. Guardians and adolescents provided informed written consent and assent, respectively. All procedures were approved by the University of Maryland Institutional Review Board (protocol #661215). The study design, hypotheses, and analytic strategy were not preregistered.

### Participants and Eligibility Criteria

Potentially interested adolescents completed an on-line screening assessment, including the abbreviated Social Phobia and Anxiety Scale for Children (SPAIC-11), which quantifies the frequency of social-anxiety symptoms across different social settings (<u>Bunnell et al., 2015</u>). Three additional *ad hoc* items (*How much does anxiety bother you or cause you distress? How much does anxiety “mess things up” for you? How much of an effect does anxiety have on your life?*) were added to the screening assessment shortly after enrollment began and were used to probe the degree of distress and impairment on a 5-point Likert scale ranging from 1 *(Not at all)* to 5 (*Extremely*). Adolescents were invited to enroll if, at the time of the preliminary screening, they showed evidence of either **(a)** *significant social anxiety*, as indexed by an abbreviated SPAIC-11 score of >15 or moderate-to-severe (≥3) distress/impairment on 1 or more of the 3 follow-up items; or **(b)** *middling-to-low social anxiety*, as indexed by an abbreviated SPAIC-11 score of <7 and, if available, little-to-no (<3) distress/impairment on the 3 additional items.

All participants and their caregivers indicated that they were right-handed, native English speakers, with a full-term pregnancy (>34 weeks) and normal or corrected-to-normal color vision. Participants reported that they were free from psychiatric medications; standard MRI contraindications; and a lifetime history of severe head injury, neurological signs and disorders, psychotic disorders, and developmental delays or disorders (e.g., autism).

In total, 78 adolescents and their caregivers participated. Twelve participants were excluded from analyses due to premature termination of the MRI assessment (*n*=3), inadequate compliance with the in-scanner rating task (<50% ratings completed; *n*=3), or insufficient usable data (*n*=6; see below for details). The final sample included 66 adolescents (*M*=15.4 years, *SD*=1.3; 60.6% female; 50.0% White, 30.3% African American/Black, 7.6% Hispanic, 6.1% Asian, 6.1% multiracial/other). The sample showed a broad spectrum of social anxiety (SPAIC: *M*=10.0, *SD*=6.0, *Range*=0.0-21.3). Half of the sample (*n*=33, 60.6% female) met the SPAIC-11 cut-off (>9.17) for probable SAD (<u>Bunnell et al., 2015</u>). The sample was significantly enriched for DSM-IV SAD diagnoses (*n*=24, 58.3% female). As expected, co-morbidity was rampant, and the majority of adolescents with SAD (*n*=14, 58.3%) received at least one other categorical diagnosis (<u>Koyuncu et al., 2019</u>; <u>Lahey et al., 2022</u>; <u>Tiego et al., 2023</u>). GAD (*n*=12, 50.0%) and MDD (*n*=8, 33.3%) were the most common co-morbidities, in broad accord with recent adolescent epidemiology studies (<u>Ernst et al., 2023</u>; <u>Jystad et al., 2021</u>).

### Clinical Assessments

#### Abbreviated Social Phobia and Anxiety Scale for Children (SPAIC-11)

Consistent with recent methodological recommendations, the 11-item SPAIC was used to quantify dimensional variation in adolescent-reported social anxiety (<u>Bunnell et al., 2015</u>; <u>Tulbure et al., 2012</u>; <u>Wong et al., 2016</u>). Prior work demonstrates that the SPAIC is reliable, sensitive to DSM diagnostic status, responsive to treatment, and correlated with independent assessments of social anxiety, including parent report and speech latency during a public-speaking challenge (<u>Beidel et al., 2000</u>; <u>Bunnell et al., 2015</u>; <u>Tulbure et al., 2012</u>; <u>Wong et al., 2016</u>). Each of the 11 items probe the frequency of fear or anxiety evoked by a specific, developmentally appropriate social situation (e.g., *“…feel scared when I have to speak or read in front of a group of people”*), using a 3-point Likert scale ranging from 0 (*never*) to 2 (*always*). Ten of the 11 items focus on subjective symptoms of fear and anxiety (*“feel scared”*), whereas the remaining item focuses on a behavioral sign (*“…do not speak to anyone until they speak to me”*). Eight of the items (e.g., *“…feel scared when I start to talk to…”*) required separate responses for different types of interaction partners (*boys or girls my age that I know, boys or girls my age that I don’t know, adults*) (<u>Beidel et al., 1995</u>). Responses for these items were averaged across partners and rounded to the nearest integer (<u>Bunnell et al., 2015</u>). Higher scores indicate more pervasive and severe social anxiety (<u>Beidel et al., 2000</u>). In the present sample, total SPAIC-11 sum scores showed acceptable reliability (α=0.94), were strongly correlated with SAD diagnostic status (*r*=0.66), and independent of age and biological sex (|*r*|<0.16). As expected, based on the observed pattern of diagnostic co-morbidity (see above), total SPAIC-11 scores were associated with the presence of other internalizing diagnoses (e.g., GAD: *r*=0.50).

#### *Mini-International* Neuropsychiatric *Interview (MINI) for Children and Adolescents*

For descriptive purposes, diagnostic status was assessed by a masters-level clinician using the Mini International Neuropsychiatric Interview for Children and Adolescents (MINI) (<u>Sheehan et al., 2010</u>). To encourage frank reporting, diagnostic interviews were completed without the caregiver present.

### Maryland Social Threat Countdown (MSTC) Paradigm

#### Task Structure

Building on prior work in adults (<u>Grogans et al., *accepted in principle*</u>; <u>Hur et al., 2022</u>; <u>Hur et al., 2020</u>; <u>Kim et al., 2023</u>), the MSTC paradigm takes the form of a 2 (*Valence:* Social Threat/Safety) × 2 (*Temporal Certainty:* Uncertain/Certain) × 2 (*Phase:* Anticipation/Presentation) repeated-measures, randomized event-related design (**Figure 1**). On Certain Social Threat trials, participants saw a descending stream of integers (‘count-down’) for 18 s. To maximize distress, the anticipation period always culminated with the presentation of a multi-modal social threat (1.8 s) encompassing a photograph of an angry young-adult face and an audio clip of a hostile verbal remark (e.g., *“No one wants you here”*). Uncertain Social Threat trials were similar, but the integer stream was randomized and presented for a temporally uncertain and variable duration (*Range*=8-30 s; *M*=18 s). On Uncertain trials, integers were randomly drawn from a near-uniform distribution ranging from 1 to 45 to reinforce the impression that Uncertain trials could be much longer than Certain ones and to minimize incidental temporal learning (‘time-keeping’). White-noise visual masks (3.2 s) were presented between trials to minimize persistence of the visual reinforcers in iconic memory. Safety trials were similar but terminated with the delivery of normatively benign photographs (happy faces) and audio clips (*“I play soccer”*). Consistent with recent recommendations (<u>Shackman & Fox, 2016</u>), the mean duration of the anticipation phase was identical across conditions, ensuring an equal number of measurements (TRs/condition). Mean duration was chosen to enhance detection of task-related differences in the blood oxygen level-dependent (BOLD) signal (<u>Henson, 2007</u>). Valence was continuously signaled throughout the anticipation epoch by the background color of the display. Certainty was signaled by the nature of the integer stream. Participants were periodically prompted (following the offset of the visual mask) to rate the intensity of fear/anxiety experienced a few moments earlier, during the anticipation period of the prior trial, using a 1-4 (*least/most*) scale and an MRI-compatible response pad (MRA, Washington, PA). Participants rated each trial type once per scan (16.7% trials). Participants were completely informed about the task design and contingencies prior to scanning. The task was administered in 3 scans, with short breaks between scans. Simulations were used to optimize the trial order and timing to minimize predictor collinearity (variance inflation factors < 1.68) (<u>Mumford et al., 2015</u>). Stimulus presentation and ratings acquisition were controlled using Presentation software (version 19.0, Neurobehavioral Systems, Berkeley, CA).

#### Procedures

Prior to fMRI scanning, participants practiced an abbreviated version of the paradigm until participants indicated and staff confirmed understanding. During scanning, foam inserts were used to immobilize the participant’s head within the head-coil and mitigate potential motion artifact. Participants were continuously monitored using an MRI-compatible eye-tracker (Eyelink 1000; SR Research, Ottawa, Ontario, Canada) and the AFNI real-time motion plugin (<u>Cox, 1996</u>). Eye-tracking data were not recorded. Measures of respiration and breathing were continuously acquired during scanning using a Siemens respiration belt and photo-plethysmograph affixed to the first digit of the non-dominant hand. Following the last scan, participants were removed from the scanner, debriefed, compensated, and discharged.

#### Visual Stimuli

A total of 72 face photographs were drawn from the Chicago Face Database (versions 1-2) (<u>Ma et al., 2015</u>). Stimuli included 36 young-adult models of varying race and sex, each depicting 1 angry and 1 happy expression. Visual stimuli were digitally back-projected onto a semi-opaque screen mounted at the head-end of the scanner bore and viewed using a mirror mounted on the head-coil (Powerlite Pro G5550, Epson America, Inc., Long Beach, CA).

#### Auditory Stimuli

A total of 72 custom auditory stimuli were created by recording 36 young-adult voice actors, recruited to match the apparent sex and race of the photograph models. Each voice actor provided 1 threatening and 1 benign audio statement, equated for the number of syllables. Voice actors were carefully coached to deliver the threatening statements (e.g., *“I don’t like you”*) in a hostile manner and to deliver the benign statements (e.g., *“Today is nice”*) in a neutral or mildly positive manner. Audio stimuli were volume standardized. To reinforce the naturalistic nature of the paradigm, each photograph was consistently paired with a specific sex- and race-matched voice actor. Auditory stimuli were delivered using an amplifier (PA-1 Whirlwind) with in-line noise-reducing filters and ear buds (S14; Sensimetrics, Gloucester, MA) fitted with noise-reducing ear plugs (Hearing Components, Inc., St. Paul, MN).

### MRI Data Acquisition

MRI data were acquired using a Siemens Magnetom TIM Trio 3T scanner (32-channel head-coil). Sagittal T1-weighted anatomical images were acquired using a magnetization prepared rapid acquisition gradient echo (MPRAGE) sequence (TR=1,900 ms; TE=2.32 ms; inversion=900 ms; flip= 9°; sagittal slice thickness=0.9 mm; in-plane=0.449 × 0.449mm; matrix=512 × 512; FOV=230 × 230). To enhance anatomical resolution, a multi-band sequence was used to collect oblique-axial echo planar imaging (EPI) volumes during the MSTC task (multiband acceleration=6; TR=1,000 ms; TE=39.4 ms; flip=36.4°; slice thickness=2.2 mm, number of slices=66; in-plane=2.1875 × 2.1875 mm; matrix=96 × 96). Images were collected in the oblique-axial plane (approximately −20° relative to the AC-PC plane) to minimize potential susceptibility artifacts. A total of three 568-volume scans were acquired. The scanner automatically discarded 7 volumes prior to the first recorded volume. To enable field map distortion correction, a pair of oblique-axial co-planar spin echo images with opposing phase encoding direction was also acquired (TR=7,220 ms; TE=73 ms; slice thickness=2.2 mm; matrix=96 × 96).

### MRI Data Pipeline

Methods were optimized to minimize spatial-normalization error and other potential sources of noise. Data were visually inspected before and after processing for quality assurance.

#### Anatomical Data Processing

Methods are similar to those described in other recent reports by our group (<u>Grogans et al., *accepted in principle*</u>; <u>Hur et al., 2018</u>; <u>Hur et al., 2022</u>; <u>Hur et al., 2020</u>; <u>Kim et al., 2023</u>). T1-weighted images were inhomogeneity corrected using *N4* (<u>Tustison et al., 2010</u>) and denoised using *ANTS* (<u>Avants et al., 2011</u>). The brain was then extracted using *BEaST* (<u>Eskildsen et al., 2012</u>) and brain-extracted and normalized reference brains from *IXI* (<u>BIAC, 2022</u>). Brain-extracted T1 images were normalized to a version of the brain-extracted 1-mm T1-weighted MNI152 (version 6) template (<u>Grabneret al., 2006</u>) modified to remove extracerebral tissue. Normalization was performed using the diffeomorphic approach implemented in *SyN* (version 2.3.4) (<u>Avants et al., 2011</u>). Brain-extracted T1 images were segmented—using native-space priors generated in *FAST* (version 6.0.4) (<u>Jenkinson et al., 2012</u>)—to enable T1-EPI co-registration (see below).

#### Fieldmap Data Processing

SE images and *topup* were used to create fieldmaps. Fieldmaps were converted to radians, median-filtered, and smoothed (2-mm). The average of the distortion-corrected SE images was inhomogeneity corrected using *N4* and masked to remove extracerebral voxels using *3dSkullStrip* (version 19.1.00).

#### Functional Data Processing

EPI files were de-spiked using *3dDespike*, slice-time corrected to the TR center using *3dTshift*, and motion corrected to the first volume and inhomogeneity corrected using *ANTS* (12-parameter affine). Transformations were saved in ITK-compatible format for subsequent use (<u>McCormick et al., 2014</u>). The first volume was extracted for EPI-T1 coregistration. The reference EPI volume was simultaneously co-registered with the corresponding T1-weighted image in native space and corrected for geometric distortions using boundary-based registration (<u>Jenkinson et al., 2012</u>). This step incorporated the previously created fieldmap, undistorted SE, T1, white matter (WM) image, and masks. The spatial transformations necessary to transform each EPI volume from native space to the reference EPI, from the reference EPI to the T1, and from the T1 to the template were concatenated and applied to the processed EPI data in a single step to minimize incidental spatial blurring. Normalized EPI data were resampled (2 mm^3^) using fifth-order b-splines. To maximize anatomical resolution, no additional spatial filters were applied, consistent with recent recommendations (<u>Tillman et al., 2018</u>) and other work by our group (<u>Grogans et al., *accepted in principle*</u>; <u>Kim et al., 2023</u>). Hypothesis testing focused on anatomically defined ROIs (see below). By convention, exploratory whole-brain voxelwise analyses employed data that were spatially smoothed (6-mm; *3DblurInMask*).

### fMRI Data Exclusions and Modeling

#### Data Exclusions

Participants who responded to <50% of rating prompts—indicating inadequate task engagement—were excluded from analyses (*n*=3). Volume-to-volume displacement (>0.5 mm) was used to assess residual motion artifact. Scans with excessively frequent artifacts (≥2.5 *SD*) were discarded. To assess task-correlated motion, we computed correlations between the design matrix and motion estimates (see above). Scans showing extreme correlations (≥ 2.5 *SD*) were discarded. Participants who failed to provide at least 2 scans of usable data were censored from analyses (*n*=6).

#### Canonical First-Level Modeling

For each participant, first-level modeling was performed using GLMs implemented in *SPM12* (version 7771), with the default autoregressive model and the temporal band-pass filter set to the hemodynamic response function (HRF) and 128 s (<u>Wellcome Centre for Human Neuroimaging, 2022</u>). Anticipatory hemodynamic signals were modeled using variable-duration rectangular (‘boxcar’) regressors spanning the anticipation (‘countdown’) epochs of the Uncertain Threat, Certain Threat, and Uncertain Safety trials; and convolved with a canonical HRF and its temporal derivative. To maximize design efficiency, Certain-Safety anticipation served as the reference condition and contributed to the baseline estimate (<u>Poline et al., 2007</u>). Epochs corresponding to the acute presentation of threatening and benign face/voice stimuli were simultaneously modeled using the same approach, separately for each of the 4 trial types (**Figure 1**). Epochs corresponding to the presentation of the white-noise visual masks and rating prompts were treated as nuisance regressors. EPI volumes acquired before the first trial and following the final trial were unmodeled and contributed to the baseline estimate. Consistent with prior work using shock-reinforced variants of the countdown task in adults (<u>Grogans et al.,</u> *<u>accepted in principle</u>;* <u>Hur et al., 2022</u>; <u>Hur et al., 2020</u>; <u>Kim et al., 2023</u>), additional nuisance variates included estimates of volume-to-volume displacement, motion (6 parameters, 0- and 1-volume lagged), cerebrospinal fluid (CSF) signal, instantaneous pulse and respiration rates, and ICA-derived nuisance signals (e.g. brain edge, CSF edge, global motion, white matter) (<u>Pruim et al., 2015</u>). Volumes with excessive volume-to-volume displacement (>0.33 mm) were censored.

#### Extended Amygdala Regions of Interest (ROIs)

Consistent with prior work by our group, task-related Ce and BST activation was quantified using well-established, anatomically defined ROIs and spatially unsmoothed fMRI data (<u>Grogans et al., *accepted in principle*</u>; <u>Hur et al., 2018</u>; <u>Kim et al., 2023</u>) (**Figure 1**). The derivation of the Ce ROI is detailed in Tillman et al. (<u>Tillman et al., 2018</u>). The probabilistic BST ROI was developed by Theiss and colleagues and thresholded at 25% (<u>Theiss et al., 2017</u>). The BST ROI mostly encompasses the supra-commissural BST, given the difficulty of reliably discriminating the borders of regions below the anterior commissure in T1-weighted MRIs (<u>Kruger et al., 2015</u>). Bilateral ROIs were decimated to the 2-mm resolution of the fMRI data. EAc ROI analyses used standardized regression coefficients extracted and averaged for each combination of task contrast (e.g., Uncertain Threat vs. Certain Safety anticipation), ROI, and participant.

### Analytic Strategy

#### Overview

The overarching goal of this study was to determine the impact of variation in social anxiety (*z*-transformed SPAIC-11) on reactivity to the MSTC paradigm (**Figure 1**). Hypothesis testing focused on two measurement modalities: **(a)** in-scanner ratings of anticipatory distress, and **(b)** and fMRI measures of EAc (Ce/BST) activation during the anticipation and presentation phases of the MSTC paradigm. Except where otherwise noted, analyses were performed using *SPSS* (Version 27.0.1). Diagnostic procedures (e.g., influence statistics) and data visualizations were used to confirm test assumptions (<u>Tukey, 1977</u>). Predictor collinearity was acceptable for all analytic models. Some figures were created using *R* (version 4.0.2), *RStudio*, and *ggplot2* (version 3.4.1) (<u>R Core Team, 2022</u>; <u>RStudio Team, 2022</u>; <u>Wickham, 2016</u>).

#### Behavioral Hypothesis Testing

A standard mixed-effects GLM was used to determine whether the anticipation of social threat elicits distress, test whether individual differences in social anxiety are associated with amplified distress, and explore the possibility that these effects are conditional on the temporal certainty of social encounters. Paralleling the MSTC paradigm, the GLM was implemented as a 2 (*Valence:* Social Threat/Safety) × 2 (*Temporal Certainty:* Uncertain/Certain) design with social anxiety (*z*-transformed SPAIC-11) included as a dimensional predictor. In effect, this model allowed us to partition variance in anticipatory distress into that associated with the experimental task, with the severity of social anxiety, and with their interaction. Significant effects were interrogated using focal contrasts (paired Students *t*-tests) and regressions as appropriate. Hotelling’s *t*-test, as implemented in *FZT* (<u>Garbin, 2024</u>), was used to compare dependent correlations (<u>Hotelling, 1940</u>).

#### fMRI Hypothesis Testing

Standard mixed-effects GLMs were used to probe the association between standardized social anxiety symptoms and EAc (Ce/BST) reactivity to the anticipation and presentation of social threat, and to explore the possibility that these effects are conditional on the temporal certainty of threat encounters. For the *anticipation* phase, the GLM was implemented as a 3 (*Condition:* Uncertain Social Threat, Certain Social Threat, and Uncertain Safety) × 2 (*Region:* Ce, BST) design with social anxiety (*z*-transformed SPAIC-11) included as a dimensional predictor. This analytic choice was dictated by the fact that the anticipation phase of Certain Safety trials served as the implicit baseline condition for first-level fMRI models (see above). Huynh-Feldt correction was used as necessary. For the *presentation* phase, the GLM was implemented as a 2 (*Valence:* Social Threat, Safety) × 2 (*Temporal Certainty:* Uncertain, Certain) × 2 (*Region:* Ce, BST) design with social anxiety again included as a dimensional predictor. For both GLMs, significant interactions were again decomposed using focal contrasts (paired Students *t*-tests) and regressions.

In head-to-head comparisons of criterion validity, simple paper-and-pencil measures often outperform more sophisticated brain imaging metrics (<u>Shackman & Fox, 2018</u>). To gauge the added explanatory value (‘incremental validity’) of statistically significant brain metrics, we re-computed the relevant multiple regressions while simultaneously incorporating paper-and-pencil distress ratings or *z*-transformed SAD diagnostic status. Sensitivity analyses confirmed that key conclusions were minimally altered by the inclusion of nuisance variation in biological sex, pubertal status (<u>Petersen et al., 1988</u>), or age (not reported).

## RESULTS

### The anticipation of social threat elicits distress

As shown in **Figure 2a**, in-scanner ratings of fearful and anxious distress were elevated during the anticipation of social threat compared to safety, and during the anticipation of temporally uncertain compared to certain social cues (*Valence: F*(1,64)=47.39, *p*<0.001, *pη^2^*=0.43; *Certainty: F*(1,64)=18.18, *p*<0.001, *pη^2^*=0.22). These observations dovetail with work focused on threat-of-shock paradigms in adults and reinforce the validity of the MSTC paradigm for probing social anxiety in adolescents (<u>Grogans et al.,</u> *<u>accepted in principle</u>;* <u>Hur et al., 2020</u>; <u>Kim et al., 2023</u>).

**Figure 2.**
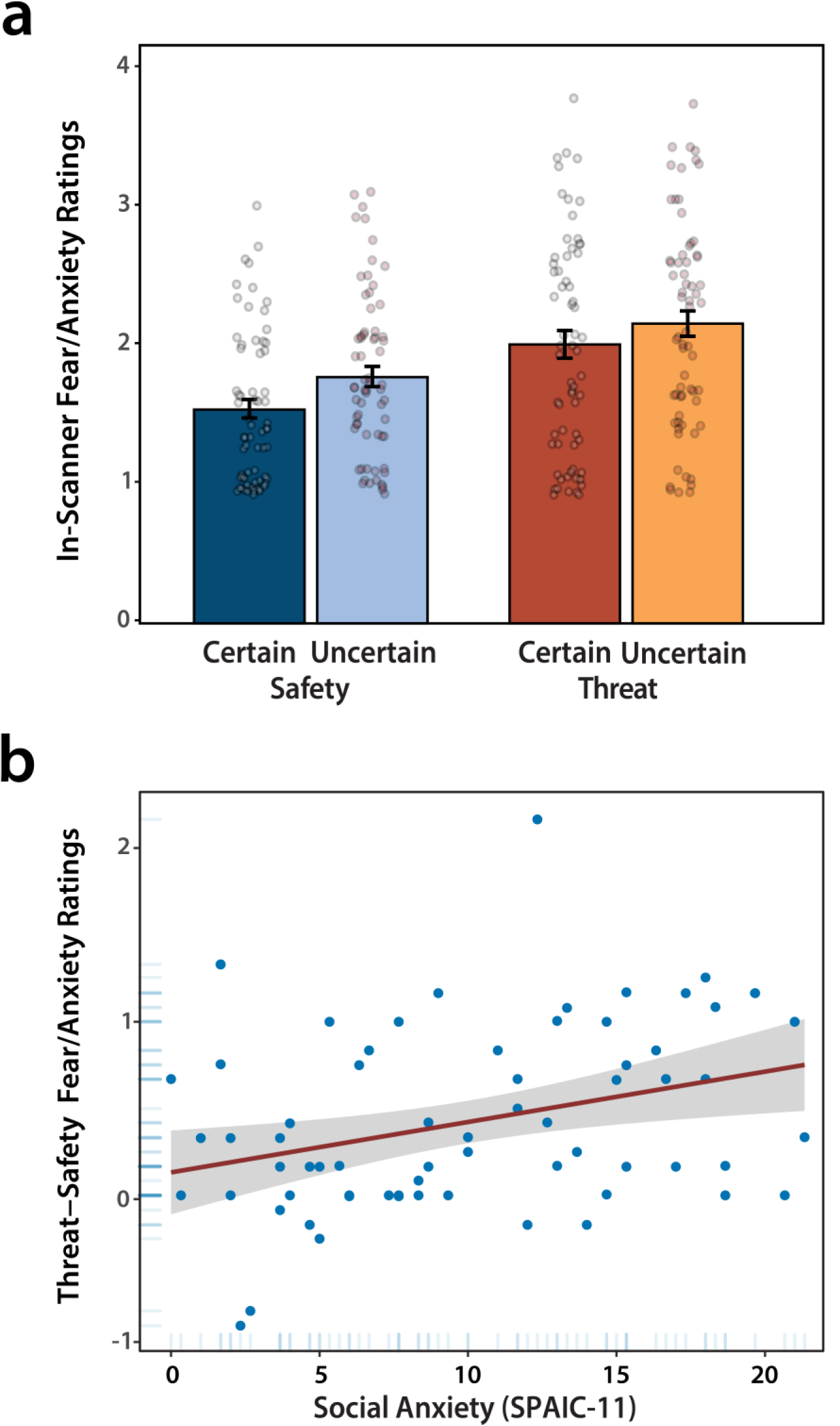
In-scanner ratings of subjective distress. **a.** Fear and anxiety were significantly elevated during the anticipation of social threat compared to safety, and during the anticipation of temporally uncertain compared to certain reinforcers (*p*<0.001). Error bars depict the standard error of the mean. Dots indicate individual participants. **b.** More anxious adolescents experience heightened distress during the anticipation of threatening, compared to benign, social cues (*p*=0.01). Red line depicts the regression slope. Gray envelope depicts the 95% confidence interval. Dots indicate individual participants.

### Social anxiety potentiates distress during the anticipation of social threat

As shown in **Figure 2b**, adolescents with more severe social anxiety reported heightened distress during the anticipation of social threat relative to safety (*Valence × Social Anxiety: F*(1,64)=7.81; *p*=0.01, *pη^2^*=0.11). Although the main effect of social anxiety was also significant (*F*(1,64)=4.80, *p*=0.03, *pη^2^*=.07), follow-up tests showed that this was entirely driven by the association with threat-elicited distress (*Threat: t*(64) = 2.79, *p* = 0.007; *Safety: t*(64) = 1.01, *p* = 0.32; *Threat-versus-Safety: tHotelling*(63) = 2.20, *p* = 0.03). Likewise, in a simultaneous regression model, only threat-related distress was significantly and uniquely associated with dimensional variation in social anxiety (*Threat: t*(63) = 2.85, *p* = 0.006; *Safety: t*(63) = −1.21, *p* = 0.23).

### The EAc is preferentially engaged by the anticipation of uncertain social threat

We used anatomically defined Ce and BST ROIs and spatially unsmoothed fMRI data to test whether more severe social anxiety symptoms are associated with amplified EAc activation when anticipating social-threat encounters, and to explore the possibility that the degree of amplification depends on the temporal certainty of encounters. A mixed-effects GLM revealed a significant effect of Region, reflecting greater activation, on average, in the Ce compared to the BST (*F*(1,64)=4.81; *p*=0.03, *pη^2^*=0.07). The Condition effect was also significant (*F*(2,128)=3.84; *p*=0.02, *pη^2^*=0.06). As shown in **Figure 3a**, the anticipation of uncertain social threat elicited greater EAc activation (averaged across regions) compared to both certain social threat and uncertain safety (*Certain Threat: t*(64)=2.56, *p*=0.01; *Uncertain Safety: t*(64)=2.07, *p*=0.04), which in turn did not significantly differ from one another (*t*(64)=-0.39, *p*=0.70). The Condition × Region interaction was not significant (*p*=0.57). Together, these results indicate that the EAc is preferentially engaged by the anticipation of uncertain social threat—consistent with prior work in adults (<u>Figel et al., 2019</u>)—but the degree of engagement does not appreciably differ across the Ce and BST, its two major divisions.

**Figure 3.**
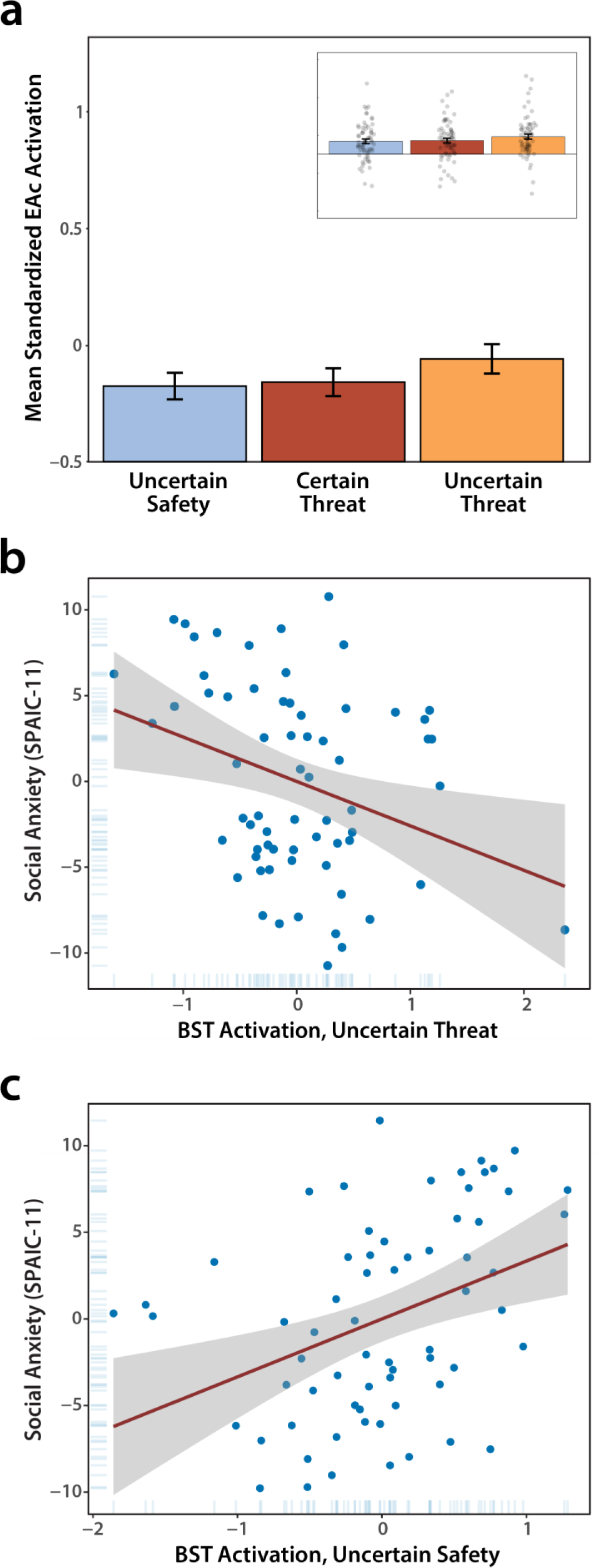
Anticipation phase of the Maryland Social Threat Countdown paradigm. **a.** The EAc is preferentially engaged by the temporally uncertain anticipation of social threat (*p*=0.03). Inset depicts individual participants. Error bars depict the standard error of the mean. **b.** Adolescents with more severe social anxiety show *diminished* BST reactivity during the temporally uncertain anticipation of social threat, while controlling for differences in Ce anticipatory activation (*p*=0.01). **c.** Socially anxious adolescents show *amplified* BST reactivity during the temporally uncertain anticipation of normatively benign social stimuli, while controlling for differences in Ce anticipatory activation (*p*=0.002). Conventions are identical to Figure 2. Abbreviations—BST, bed nucleus of the stria terminalis; EAc, central extended amygdala (Ce/BST).

### Social anxiety is associated with diminished BST discrimination of anticipated threat and safety

The GLM also revealed significant Condition × Social Anxiety (*F*(2,128)=5.87, *p*=0.004, *pη^2^*=0.08) and Condition × Region × Social Anxiety interactions (*F*(2,128)=4.75; *p*=0.01, *pη^2^*=0.07). Other effects were not significant (*p*>0.46). To decompose the significant three-way interaction, we computed separate regression models for the Ce and BST, in each case including reactivity to each of the three anticipation conditions—uncertain social threat, certain social threat, and uncertain safety—as simultaneous predictors of social anxiety. Results indicated that Ce activation is unrelated to variation in social anxiety (|*t*|(62)<1.19, *p*>0.23). On the other hand, *blunted* BST reactivity to uncertain-threat anticipation and *heightened* BST reactivity to uncertain-safety anticipation were both associated with more severe social anxiety (*Uncertain Threat: t*(62)=-2.52, *p*=0.01; *Uncertain Safety: t*(62)=2.34, *p*=0.02), with negligible effects evident for certain-threat anticipation (*t*(62)=0.17, *p*=0.87). As shown in **Figures 3b-c**, both associations remained significant while controlling for variation in Ce activation, indicating a statistically unique association between BST reactivity and the severity of adolescent social anxiety (*Uncertain Threat: t*(60)=-2.57, *p*=0.01; *Uncertain Safety: t*(60)=3.23, *p*=0.002).

These results demonstrate that the most robust EAc activation occurs during the uncertain anticipation of social threat, with significantly lower activation evident during the uncertain anticipation of encounters with normatively safe social cues (**Figure 3a**). This suggests that the normative mean difference across the two conditions is diminished in the BST of adolescents with more severe social anxiety. Consistent with this intuition, higher social anxiety was negatively associated with the simple arithmetic difference in BST activation across the two conditions (*Uncertain Social Threat – Uncertain Safety, anticipation: r*(64)=-0.38 [95% CI: 0.15, 0.57], *p*=0.002). Although exploratory voxelwise analyses revealed a number of regions that were recruited during social-threat anticipation, they uncovered no whole-brain significant associations with social anxiety (**Supplementary Tables S1-S7**).

To clarify the explanatory value of BST reactivity, we computed a new regression model, using a combination of the BST neuroimaging metrics and two self-report metrics acquired during the MSTC— indiscriminate distress (Threat + Safety Ratings) and threat-potentiated distress (Threat – Safety Ratings)—to predict social anxiety (cf. **Figure 2**). Results indicated that BST activation during the anticipation of uncertain threat and safety continued to explain significant variance in social anxiety while controlling for both forms of task-related distress (*Uncertain Threat: t*(61)=-2.85, *p*=.006; *Uncertain Safety: t*(61)=2.86, *p*=.006; *Ratings: t*(61)<1.48, *p*>.14). The same conclusion was also evident using the difference in anticipatory BST activation across the two conditions (*t*(62)=-2.76, *p*=0.008) or when controlling for all four condition-specific distress ratings (|*t*|(59)>2.23, *p*<0.03). The difference in anticipatory BST activation also explained significant variance in the severity of social anxiety while controlling for SAD diagnostic status (*t*(62)=-2.62, *p*=0.01). In short, BST anticipatory activation explains dimensional variation in adolescent social anxiety above-and-beyond that accounted for by more traditional self- and clinician-assessments, underscoring the added value (‘incremental validity’) of the fMRI metrics. These observations also make it clear that the association between indiscriminate BST anticipatory activation and social anxiety cannot be attributed to simple differences in subjective reactivity to the MSTC paradigm.

### The EAc is robustly recruited by the presentation of social threat, independent of social anxiety

As shown in **Figure 4**, a mixed-effects GLM indicated that the EAc (Ce and BST) was more strongly engaged by the presentation of threatening compared to normatively benign faces and voices (*Valence: F*(1,64)=8.03; *p*=0.006, *pη^2^*=0.11). Results also demonstrated that the Ce was more responsive to the acute presentation of social cues—irrespective of their valence—when compared to the BST (*Region: F*(1,64)=11.56; *p*=0.001, *pη^2^*=0.15). No other effects were significant (*p*>0.18). In sum, while the EAc, in aggregate, is robustly recruited by the acute presentation of threatening face-voice stimuli, this is largely independent of variation in the severity of social anxiety symptoms.

**Figure 4.**
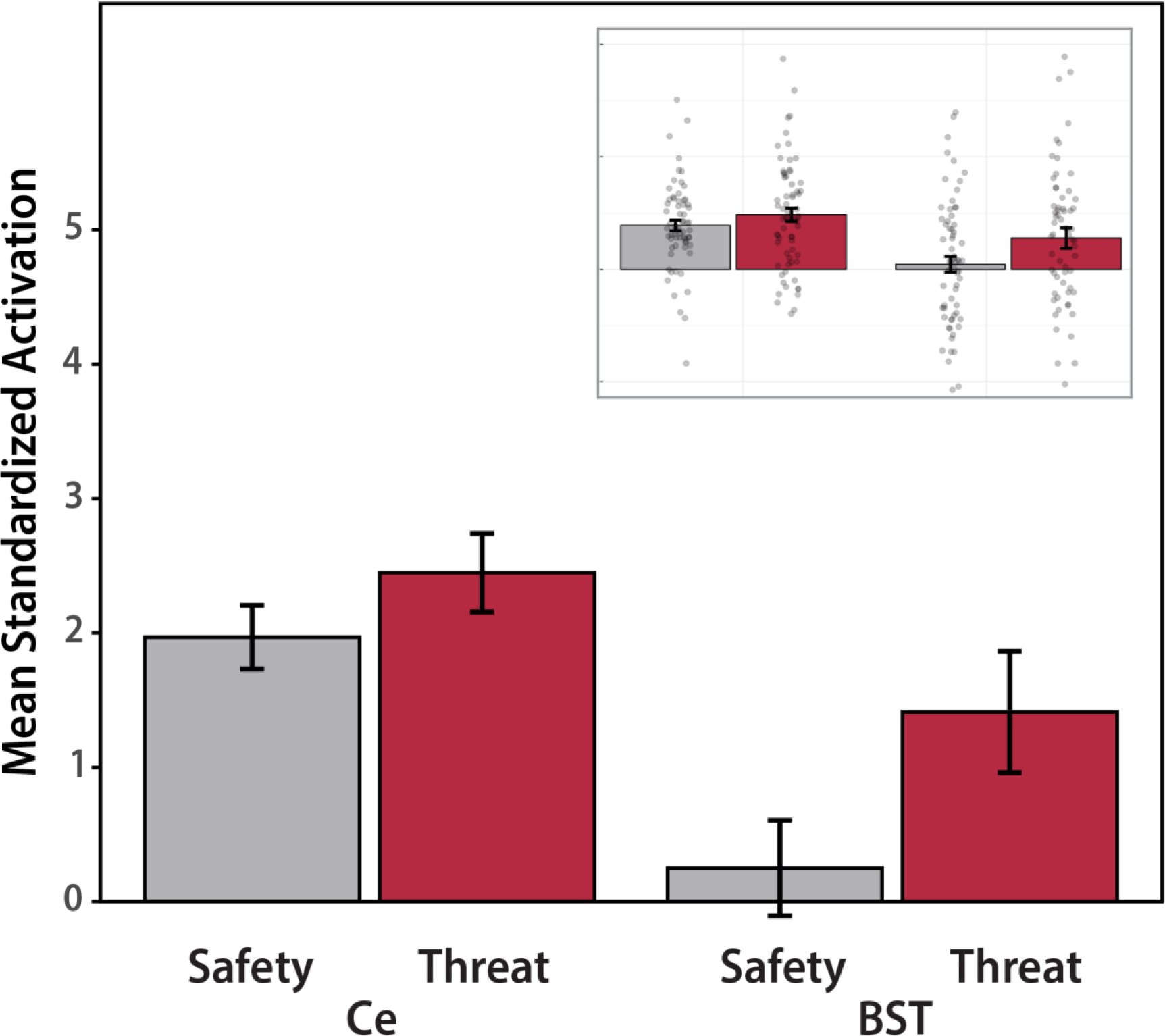
Presentation phase of the Maryland Social Threat Countdown paradigm. GLM results demonstrated that the EAc was more strongly recruited by the acute presentation of threatening (relative to benign) social stimuli (*p*=0.006) and that the Ce is more responsive (relative to the BST) to faces and voices, irrespective of their valence (*p*=0.001). Inset depicts individual participants. Other conventions are identical to Figure 2. Abbreviations—BST, bed nucleus of the stria terminalis.

As expected, a whole-brain voxelwise analysis confirmed that the presentation of faces and voices recruited the amygdala and relevant areas of sensory cortex, including the bilateral fusiform gyri, superior temporal sulci, and Heschl’s gyri (**Supplementary Tables S8-S10**). Exploratory analyses did not detect any whole-brain significant voxelwise associations with dimensional variation in social anxiety (**Supplementary Tables S11-22**).

## DISCUSSION

Among socially anxious adolescents, distress and impairment can be triggered by both acute social scrutiny, such as interacting with unfamiliar peers, and settings where negative evaluations are possible, but uncertain in their timing or likelihood, as when first entering a public restroom or navigating a school hallway. To date, comparatively little scientific attention has been devoted to understanding the neural systems underlying heightened reactivity to anticipated social threat. Here we leveraged a novel fMRI paradigm—the MSTC task—in a racially diverse sample of adolescents selectively recruited to encompass a broad spectrum of social anxiety symptoms (50% adolescents of color; 50.0% positive SAD screen; 36.4% SAD diagnosis) (**Figure 1**). Results demonstrated that fearful and anxious feelings were significantly elevated, on average, during the anticipation of threatening social cues, confirming the basic validity of the MSTC task as a probe of social anxiety (**Figure 2a**). Adolescents with more severe social anxiety reported potentiated distress when anticipating encounters with social threat (**Figure 2b**). Neuroimaging results revealed robust EAc (Ce and BST) activation during the uncertain anticipation of social threat, relative to the anticipation of both certain threat and uncertain safety (**Figure 3a**). Adolescents with more severe social anxiety showed *blunted* BST activation during the uncertain anticipation of social threat and *heightened* BST activation during the uncertain anticipation of benign social cues, indicating diminished discrimination of the two contexts. Indeed, follow-up analyses demonstrated that more severe symptoms were also associated with a reduction in the difference in BST activation across the two conditions. These brain-anxiety associations remained significant when controlling for Ce reactivity, paper-and-pencil measures of task-related distress, or SAD diagnostic status, underscoring the unique explanatory value of BST anticipatory activity (**Figures 3b-c**). Although the EAc (Ce and BST) was robustly engaged during the acute presentation of threatening faces and voices—consistent with preclinical research in adults (<u>Sladky et al., 2018</u>)—variation in the degree of engagement was unrelated to the severity of social anxiety symptoms (**Figure 4**).

During the anticipation phase of the MSTC paradigm, the EAc (Ce and BST) was much more strongly recruited, on average, by uncertain social threat compared to either certain threat or uncertain safety. Within the BST (but not the Ce), these normative mean differences are further moderated by individual differences in the severity of social anxiety symptoms. Adolescents with more severe symptoms show blunted BST activation during the uncertain anticipation of hostile social cues, and amplified activation during the uncertain anticipation of benign ones. Might this pattern of results simply reflect heightened BST reactivity to benign social stimuli (**Figure 1**)? After all, socially anxious adolescents and adults often experience heightened levels of dread and discomfort during the anticipation or experience of positive social interactions (e.g., receiving praise) (<u>Fredrick & Luebbe, 2020</u>; <u>Weeks et al., 2019</u>), perceive positive and neutral facial expressions as less approachable and more arousing (<u>Kivity & Huppert, 2015</u>), and in some cases have been shown to exhibit exaggerated amygdala activation to positive and neutral faces (<u>Birbaumer et al., 1998</u>; <u>Cooney et al., 2006</u>; <u>Crane et al., 2021</u>; <u>Straube et al., 2005</u>). The answer appears to be ‘no,’ insofar as our results demonstrate that social anxiety was unrelated to EAc activation during the acute presentation of social cues. When viewing faces and voices, the EAc is more responsive to threatening than benign cues, independent of social anxiety. This observation is consistent with work in adults with SAD (<u>Ziv et al., 2013</u>) and contraindicates a straightforward ‘hyper-reactivity to benign social cues’ explanation. Variation in the severity of social anxiety was also unrelated to EAc activation during the certain anticipation of threat, contraindicating the possibility that socially anxious adolescents are simply more pessimistic in their expectations or nonspecifically hyper-reactive to anticipated social encounters. What mechanism, then, might explain this pattern of results? Grupe and Nitschke’s conceptual framework argues that **(a)** uncertainty amplifies the salience of anticipated encounters with threat; and **(b)** anxious individuals have difficulty discriminating threat from safety (<u>Grupe & Nitschke, 2013</u>). Such biases tend to manifest most clearly in ‘weak’ situations, where there is more opportunity for individual differences in anxiety to guide thoughts, feelings, and behavior. This framework suggests that among adolescents with more severe social anxiety symptoms, the amplifying consequences of temporal uncertainty on BST activation are especially potent when anticipating weakly or ambiguously threatening social cues (unfamiliar adults emitting neutral or mildly positive expressions); comparatively weaker when anticipating more intense and normatively threatening social cues (unfamiliar adults emitting overtly hostile expressions); and absent during the certain anticipation or acute presentation of social cues. This perspective accounts for the overall pattern of mean differences across both phases of the MSTC paradigm and explains the opposing consequences of individual differences in social anxiety on BST anticipatory activity. More broadly, these observations reinforce the hypothesis that alterations in BST function underlie socially anxious adolescents’ maladaptive responses to benign-but-uncertain everyday social encounters, such as anticipating an interaction with a sales associate or asking a routine question in class.

Clearly, a number of important challenges remain for future research. First, our study was focused on a racially diverse sample of adolescents. Moving forward, it will be useful to expand this to encompass larger and more demographically representative samples. Larger samples will also afford that statistical power necessary to tease apart whether the effects that we ascribe to social anxiety actually reflect associations with broader psychopathology dimensions, such as the fear disorders or the internalizing spectrum (<u>Conway et al., 2019</u>; <u>Tiego et al., 2023</u>; <u>Watson et al., 2022</u>). Second, our analyses relied on adolescent-reported social anxiety symptoms. While this approach is common in both research and clinical settings, recent work highlights the value of using more sophisticated multi-informant approaches to quantify the depth and breadth of social anxiety (<u>Beidel et al., 2019</u>; <u>De Los Reyes et al., 2023</u>; <u>Makol et al., 2020</u>). Third, our cross-sectional design does not license mechanistic claims. Prospective-longitudinal and intervention research will be necessary to determine whether diminished discrimination of uncertain social threat and safety in the BST forecasts the emergence of clinically significant distress and impairment, and whether this bias is normalized by clinically effective treatments. Preliminary work in rodents is encouraging (<u>Bruzsik et al., 2021</u>; <u>De Bundel et al., 2016</u>; <u>Duvarci et al., 2009</u>), demonstrating that the BST plays a critical role in discriminating Pavlovian threat from safety. For example, blockade of BST dopamine D2 receptor signaling has been shown to potentiate defensive responses to safety cues (CS-) and blunt defensive responses to threat cues (CS+); in contrast, stimulation of this pathway enhances the discrimination of threat from safety (<u>De Bundel et al., 2016</u>). Moving forward, it will be helpful to determine whether these perturbations produce parallel changes in defensive responding to uncertain social threat (e.g., novel conspecific) and safety (e.g., novel testing chamber). Fourth, although our results highlight the importance of the BST, social anxiety is a complex phenotype that likely reflects multiple distributed networks. It will be important to understand how interactions between the BST and other brain regions support variation in specific facets of pediatric social anxiety.

In conclusion, the present results demonstrate that adolescents with more severe social anxiety symptoms show reduced discrimination of uncertain social threat and safety in the BST. These discoveries provide a neuroscientifically grounded framework for conceptualizing adolescent social anxiety and set the stage for the kinds of targeted follow-up research that will be necessary to clarify causation and, ultimately, to develop more effective or tolerable biological interventions for youth living with social anxiety. A diverse sample enriched for clinically significant levels of social anxiety, a well-controlled task, and a statistically unbiased (anatomical ROI) analytic approach enhance confidence in the robustness and translational relevance of these results.

## Supporting information

Supplementary Results

## ACKNOWLEDGMENTS

Authors acknowledge assistance and critical feedback from two anonymous reviewers, J. Blanchard, L. Dougherty, N. Fox, L. Friedman, S. Grogans, C. Kaplan, B. Nacewicz, L. Pessoa, T. Riggins, B. Stephenson, Z. Tillman, members of the Affective and Translational Neuroscience laboratory and Comprehensive Assessment and Intervention Program, and the staff of the Maryland Neuroimaging Center. This work was partially supported by the National Institutes of Health (AA030042, DA040717, MH107444, MH121409, MH131264); National Research Foundation of Korea (2021R1F1A106338513 and 2021S1A5A2A0307022913); and Yonsei Signature Research Cluster Program (2021-22-0005). All procedures were approved by the University of Maryland, College Park Institutional Review Board (protocol #661215). Authors declare no conflicts of interest. A preprint of this report is available at *bioRxiv* (https://www.biorxiv.org/content/10.1101/2023.10.30.564701v1).

## AUTHOR CONTRIBUTIONS

R.M.T., A.J.S., and J.F.S. designed the overall study with guidance from A.D.L.R. and K.A.D. R.M.T., M.D.S., and J.F.S. developed and optimized the imaging paradigm. R.M.T. managed data collection, clinical assessment, and study administration with guidance from K.A.D. R.M.T., H.C.K., A.S.A., S.I., and M.D.S. collected data. J.F.S. developed data processing and analytic software for imaging analyses. J.F.S. and R.M.T. processed imaging data. R.M.T, J.F.S., J.H., and A.J.S. analyzed data. R.M.T. and A.J.S. developed the analytic strategy. R.M.T., J.H., A.D.L.R., and A.J.S. interpreted data. R.M.T., A.J.S., J.H., and M.D.S. wrote the paper. P.D., H.C.K., J.H., and R.M.T. created figures and tables. J.H. coordinated editorial communications. A.J.S. funded and supervised all aspects of the study. All authors contributed to reviewing and revising the paper and approved the final version.

## RESOURCE SHARING

Study materials, processed data, and statistical code are publicly available (https://osf.io/aj64g/). Key neuroimaging maps are publicly available (https://neurovault.org/collections/15620). Processed neuroimaging data have also been shared with the ENIGMA Consortium Anxiety Workgroup (https://enigma.ini.usc.edu/ongoing/enigma-anxiety).

## GENERAL SCIENTIFIC SUMMARY

Clinically significant levels of social anxiety often emerge in adolescence and existing treatments are far from curative for many, underscoring the urgency of developing a deeper understanding of the underlying neurobiology. Leveraging a racially diverse sample of adolescents and a novel brain imaging paradigm, the present results show that more severe social anxiety is associated with diminished discrimination of anticipated social threat and safety in the bed nucleus of the stria terminalis (BST), a brain region thought to play a key role in promoting pathological fear and anxiety. These observations reinforce the hypothesis that alterations in BST function underlie socially anxious adolescents’ maladaptive responses to benign-but-uncertain everyday social encounters—such as anticipating an interaction with a sales associate or asking a routine question in class—and set the stage for targeted replication and extension studies.

## REFERENCES

Angst, J., Paksarian, D., Cui, L., Merikangas, K. R., Hengartner, M. P., Ajdacic-Gross, V., & Rössler, W. (2016). The epidemiology of common mental disorders from age 20 to 50: results from the prospective Zurich cohort Study. Epidemiology and psychiatric sciences, 25, 24–32. 10.1017/S204579601500027X

APA. (2022). Diagnostic and statistical manual of mental disorders, text revision (DSM-5-TR) (5 ed.). American Psychiatric Publishing.

Avants, B. B., Tustison, N. J., Song, G., Cook, P. A., Klein, A., & Gee, J. C. (2011). A reproducible evaluation of ANTs similarity metric performance in brain image registration [Article]. Neuroimage, 54, 2033–2044. 10.1016/j.neuroimage.2010.09.025

Batelaan, N. M., Bosman, R. C., Muntingh, A., Scholten, W. D., Huijbregts, K. M., & van Balkom, A. (2017). Risk of relapse after antidepressant discontinuation in anxiety disorders, obsessive-compulsive disorder, and post-traumatic stress disorder: systematic review and meta-analysis of relapse prevention trials. BMJ, 358, j3927. 10.1136/bmj.j3927

Bechara, A., Tranel, D., Damasio, H., Adolphs, R., Rockland, C., & Damasio, A. R. (1995). Double dissociation of conditioning and declarative knowledge relative to the amygdala and hippocampus in humans. Science, 269, 1115–1118.

Beesdo-Baum, K., & Knappe, S. (2012). Developmental epidemiology of anxiety disorders. Child Adolesc Psychiatr Clin N Am, 21(3), 457–478. 10.1016/j.chc.2012.05.001

Beidel, D., Le, T.-A. P., & Willis, E. (2019). Social anxiety disorder: An update on diagnostics, epidemiology, etiology, assessment, treatment, unanswered questions, and future directions. In S. N. Compton, M. A. Villabø, & H. Kristensen (Eds.), Pediatric anxiety disorders (pp. 201–223). Academic Press. 10.1016/B978-0-12-813004-9.00010-4

Beidel, D. C., Turner, S. M., Hamlin, K., & Morris, T. L. (2000). The Social Phobia and Anxiety Inventory for Children (SPAI–C): External and discriminative validity. Behavior Therapy, 31, 75–87. 10.1016/S0005-7894(00)80005-2

Beidel, D. C., Turner, S. M., & Morris, T. L. (1995). A new inventory to assess childhood social anxiety and phobia: The Social Phobia and Anxiety Inventory for Children. Psychological Assessment, 7, 73–79. 10.1037/1040-3590.7.1.73

BIAC. (2022). IXI Dataset. Imperial College London. Retrieved April 19 from https://brain-development.org/ixi-dataset/

Birbaumer, N., Grodd, W., Diedrich, O., Klose, U., Erb, M., Lotze, M., … Flor, H. (1998). fMRI reveals amygdala activation to human faces in social phobics. Neuroreport, 9(6), 1223–1226. 10.1097/00001756-199804200-00048

Boehme, S., Ritter, V., Tefikow, S., Stangier, U., Strauss, B., Miltner, W. H., & Straube, T. (2014). Brain activation during anticipatory anxiety in social anxiety disorder. Soc Cogn Affect Neurosci, 9, 1413–1418. 10.1093/scan/nst129

Brühl, A. B., Delsignore, A., Komossa, K., & Weidt, S. (2014). Neuroimaging in social anxiety disorder—a meta-analytic review resulting in a new neurofunctional model. Neurosci Biobehav Rev, 47, 260–280. 10.1016/j.neubiorev.2014.08.003

Bruzsik, B., Biro, L., Zelena, D., Sipos, E., Szebik, H., Sarosdi, K. R., … Toth, M. (2021). Somatostatin neurons of the bed nucleus of stria terminalis enhance associative fear memory consolidation in mice. J Neurosci, 41, 1982–1995. 10.1523/jneurosci.1944-20.2020

Bunnell, B. E., Beidel, D. C., Liu, L., Joseph, D. L., & Higa-McMillan, C. (2015). The SPAIC-11 and SPAICP-11: Two brief child- and parent-rated measures of social anxiety. J Anxiety Disord, 36, 103–109. 10.1016/j.janxdis.2015.10.002

Cain, C. K. (2023). Beyond fear, extinction, and freezing: Strategies for improving the translational value of animal conditioning research. Curr Top Behav Neurosci, 64, 19–57. 10.1007/7854_2023_434

Chavanne, A. V., & Robinson, O. J. (2021). The overlapping neurobiology of adaptive and pathological anxiety: a meta-analysis of functional neural activation. American Journal of Psychiatry, 178, 156–164. 10.1176/appi.ajp.2020.19111153

Chen, W. H., Lien, C. C., & Chen, C. C. (2022). Neuronal basis for pain-like and anxiety-like behaviors in the central nucleus of the amygdala. Pain, 163(3), e463–e475. 10.1097/j.pain.0000000000002389

Clauss, J. A., & Blackford, J. U. (2012). Behavioral inhibition and risk for developing social anxiety disorder: a meta-analytic study. J Am Acad Child Adolesc Psychiatry, 51, 1066–1075 S0890-8567(12)00592-8 [pii] 10.1016/j.jaac.2012.08.002

Conway, C. C., Forbes, M. K., Forbush, K. T., Fried, E. I., Hallquist, M. N., Kotov, R., … Eaton, N.R.. (2019). A Hierarchical Taxonomy of Psychopathology can reform mental health research. Perspectives on Psychological Science, 14, 419–436.

Cooney, R. E., Atlas, L. Y., Joormann, J., Eugène, F., & Gotlib, I. H. (2006). Amygdala activation in the processing of neutral faces in social anxiety disorder: is neutral really neutral? Psychiatry Res, 148(1), 55–59. 10.1016/j.pscychresns.2006.05.003

Cox, R. W. (1996). AFNI: Software for analysis and visualization of functional magnetic resonance neuroimages. Computers and Biomedical Research, 29, 162–173.

Crane, N. A., Chang, F., Kinney, K. L., & Klumpp, H. (2021). Individual differences in striatal and amygdala response to emotional faces are related to symptom severity in social anxiety disorder. Neuroimage Clin, 30, 102615. 10.1016/j.nicl.2021.102615

Cuijpers, P., Miguel, C., Ciharova, M., Harrer, M., Basic, D., Cristea, I. A., … Karyotaki, E. (2024). Absolute and relative outcomes of psychotherapies for eight mental disorders: a systematic review and meta-analysis. World Psychiatry, 23, 267–275. 10.1002/wps.21203

Davies, C. D., Young, K., Torre, J. B., Burklund, L. J., Goldin, P. R., Brown, L. A., … Craske, M.G.. (2017). Altered time course of amygdala activation during speech anticipation in social anxiety disorder. J Affect Disord, 209, 23–29. 10.1016/j.jad.2016.11.014

Davis, M., Walker, D. L., Miles, L., & Grillon, C. (2010). Phasic vs sustained fear in rats and humans: Role of the extended amygdala in fear vs anxiety. Neuropsychopharmacology, 35, 105–135. npp2009109 [pii] 10.1038/npp.2009.109

Davis, M., & Whalen, P. J. (2001). The amygdala: vigilance and emotion. Mol Psychiatry, 6, 13–34.

De Bundel, D., Zussy, C., Espallergues, J., Gerfen, C. R., Girault, J. A., & Valjent, E. (2016). Dopamine D2 receptors gate generalization of conditioned threat responses through mTORC1 signaling in the extended amygdala. Mol Psychiatry, 21, 1545–1553. 10.1038/mp.2015.210

De Los Reyes, A., Wang, M., Lerner, M. D., Makol, B. A., Fitzpatrick, O. M., & Weisz, J. R. (2023). The Operations Triad Model and youth mental health assessments: Catalyzing a paradigm shift in measurement validation. Journal of Clinical Child & Adolescent Psychology, 52, 19–54. 10.1080/15374416.2022.2111684

Duvarci, S., Bauer, E. P., & Paré, D. (2009). The bed nucleus of the stria terminalis mediates inter-individual variations in anxiety and fear. J Neurosci, 29, 10357–10361. 29/33/10357 [pii] 10.1523/JNEUROSCI.2119-09.2009

Emery, N. J., Capitanio, J. P., Mason, W. A., Machado, C. J., Mendoza, S. P., & Amaral, D. G. (2001). The effects of bilateral lesions of the amygdala on dyadic social interactions in rhesus monkeys (Macacamulatta). Behavioral neuroscience, 115, 515–544. http://www.ncbi.nlm.nih.gov/entrez/query.fcgi?cmd=Retrieve&db=PubMed&dopt=Citation&list_uids=11439444

Ernst, J., Ollmann, T. M., König, E., Pieper, L., Voss, C., Hoyer, J., … Beesdo-Baum, K., (2023). Social anxiety in adolescents and young adults from the general population: an epidemiological characterization of fear and avoidance in different social situations. Current Psychology, 42, 28130–28145. 10.1007/s12144-022-03755-y

Eskildsen, S. F., Coupé, P., Fonov, V., Manjón, J. V., Leung, K. K., Guizard, N., … Alzheimer’s Disease Neuroimaging Initiative. (2012). BEaST: brain extraction based on nonlocal segmentation technique. Neuroimage, 59, 2362–2373.

Evans, R., Clark, D. M., & Leigh, E. (2021). Are young people with primary social anxiety disorder less likely to recover following generic CBT compared to young people with other primary anxiety disorders? A systematic review and meta-analysis. Behav Cogn Psychother, 49(3), 352–369. 10.1017/s135246582000079x

Fehm, L., Beesdo, K., Jacobi, F., & Fiedler, A. (2008). Social anxiety disorder above and below the diagnostic threshold: prevalence, comorbidity and impairment in the general population. Soc Psychiatry Psychiatr Epidemiol, 43, 257–265. 10.1007/s00127-007-0299-4

Feinstein, J. S., Adolphs, R., Damasio, A., & Tranel, D. (2011). The human amygdala and the induction and experience of fear. Current Biology, 21, 1–5.

Feinstein, J. S., Adolphs, R., & Tranel, D. (2016). A tale of survival from the world of Patient S.M. In D. G. Amaral & R. Adolphs (Eds.), Living without an amygdala. Guilford.

Figel, B., Brinkmann, L., Buff, C., Heitmann, C. Y., Hofmann, D., Bruchmann, M., … Straube, T. (2019). Phasic amygdala and BNST activation during the anticipation of temporally unpredictable social observation in social anxiety disorder patients. Neuroimage Clin, 22, 101735. 10.1016/j.nicl.2019.101735

Fox, A. S., & Kalin, N. H. (2014). A translational neuroscience approach to understanding the development of social anxiety disorder and its pathophysiology. American Journal of Psychiatry, 171, 1162–1173.

Fox, A. S., Oler, J. A., Shackman, A. J., Shelton, S. E., Raveendran, M., McKay, D. R., … Kalin, N.H.. (2015). Intergenerational neural mediators of early-life anxious temperament. Proceedings of the National Academy of Sciences USA, 112, 9118–9122.

Fox, A. S., Oler, J. A., Tromp, D. P., Fudge, J. L., & Kalin, N. H. (2015). Extending the amygdala in theories of threat processing. Trends Neurosci, 38, 319–329. 10.1016/j.tins.2015.03.002

Fox, A. S., & Shackman, A. J. (2019). The central extended amygdala in fear and anxiety: Closing the gap between mechanistic and neuroimaging research. Neuroscience letters, 693, 58–67. 10.1016/j.neulet.2017.11.056

Fredrick, J. W., & Luebbe, A. M. (2020). Fear of positive evaluation and social anxiety: A systematic review of trait-based findings. J Affect Disord, 265, 157–168. 10.1016/j.jad.2020.01.042

Garbin, C. P. (2024). FZT. Department of Psychology. Retrieved April 21 from https://psych.unl.edu/psycrs/statpage/comp.html

Gentili, C., Cristea, I. A., Angstadt, M., Klumpp, H., Tozzi, L., Phan, K. L., & Pietrini, P. (2016). Beyond emotions: A meta-analysis of neural response within face processing system in social anxiety. Exp Biol Med (Maywood), 241(3), 225–237. 10.1177/1535370215603514

Grabner, G., Janke, A. L., Budge, M. M., Smith, D., Pruessner, J., & Collins, D. L. (2006). Symmetric atlasing and model based segmentation: an application to the hippocampus in older adults. Med Image Comput Comput Assist Interv Int Conf Med Image Comput Comput Assist Interv, 9, 58–66.

Gregory, A. M., Caspi, A., Moffitt, T. E., Koenen, K., Eley, T. C., & Poulton, R. (2007). Juvenile mental health histories of adults with anxiety disorders. Am J Psychiatry, 164, 301–308. 164/2/301 [pii] 10.1176/appi.ajp.164.2.301

Grogans, S. E., Hur, J., Barstead, M. G., Anderson, A. S., Islam, S., Kuhn, M., … Shackman, A.J.. (*accepted in principle*). Neuroticism/negative emotionality is associated with increased reactivity to uncertain threat in the bed nucleus of the stria terminalis, not the amygdala. Journal of Neuroscience [preprint available at BioRxiv].

Grupe, D. W., & Nitschke, J. B. (2013). Uncertainty and anticipation in anxiety: an integrated neurobiological and psychological perspective. Nat Rev Neurosci, 14, 488–501. nrn3524 [pii] 10.1038/nrn3524

Henson, R. (2007). Efficient experimental design for fMRI. In K. Friston, J. Ashburner, S. Kiebel, T. Nichols, & W. Penny (Eds.), Statistical Parametric Mapping: The Analysis of Functional Brain Images (pp. 193–210). Academic Press.

Hotelling, H. (1940). The selection of variates for use in prediction with some comments on the general problem of nuisance parameters. Annals of Mathematical Statistics, 11, 271–283.

Hur, J., Kaplan, C. M., Smith, J. F., Bradford, D. E., Fox, A. S., Curtin, J. J., & Shackman, A. J. (2018). Acute alcohol administration dampens central extended amygdala reactivity. Scientific Reports, 8, 16702.

Hur, J., Kuhn, M., Grogans, S. E., Anderson, A. S., Islam, S., Kim, H. C., … Shackman, A.J.. (2022). Anxiety-related frontocortical activity is associated with dampened stressor reactivity in the real world. Psychological Science, 33, 906–924. 10.1101/2021.03.17.435791

Hur, J., Smith, J. F., DeYoung, K. A., Anderson, A. S., Kuang, J., Kim, H. C., … Shackman, A.J.. (2020). Anxiety and the neurobiology of temporally uncertain threat anticipation. Journal of Neuroscience, 40, 7949–7964. 10.1101/2020.02.25.964734

Hur, J., Stockbridge, M. D., Fox, A. S., & Shackman, A. J. (2019). Dispositional negativity, cognition, and anxiety disorders: An integrative translational neuroscience framework. Progress in Brain Research, 247, 375–436.

Hyett, M. P., & McEvoy, P. M. (2018). Social anxiety disorder: looking back and moving forward. Psychol Med, 48, 1937–1944. 10.1017/s0033291717003816

Inman, C. S., Bijanki, K. R., Bass, D. I., Gross, R. E., Hamann, S., & Willie, J. T. (2020). Human amygdala stimulation effects on emotion physiology and emotional experience. Neuropsychologia, 145, 106722. 10.1016/j.neuropsychologia.2018.03.019

James, A. C., Reardon, T., Soler, A., James, G., & Creswell, C. (2020). Cognitive behavioural therapy for anxiety disorders in children and adolescents. Cochrane Database Syst Rev, 11, Cd013162. 10.1002/14651858.CD013162.pub2

Jarcho, J. M., Leibenluft, E., Walker, O. L., Fox, N. A., Pine, D. S., & Nelson, E. E. (2013). Neuroimaging studies of pediatric social anxiety: paradigms, pitfalls and a new direction for investigating the neural mechanisms. Biol Mood Anxiety Disord, 3, 14. 10.1186/2045-5380-3-14

Jenkinson, M., Beckmann, C. F., Behrens, T. E., Woolrich, M. W., & Smith, S. M. (2012). FSL. Neuroimage, 62, 782–790. 10.1016/j.neuroimage.2011.09.015

Jystad, I., Bjerkeset, O., Haugan, T., Sund, E. R., & Vaag, J. (2021). Sociodemographic correlates and mental health comorbidities in adolescents with social anxiety: The Young-HUNT3 Study, Norway. Front Psychol, 12, 663161. 10.3389/fpsyg.2021.663161

Kalin, N. H., Fox, A. S., Kovner, R., Riedel, M. K., Fekete, E. M., Roseboom, P. H., … Oler, J.A.. (2016). Overexpressing corticotropin-releasing hormone in the primate amygdala increases anxious temperament and alters its neural circuit. Biol Psychiatry, 80, 345–355. 10.1016/j.biopsych.2016.01.010

Kalin, N. H., Shelton, S. E., & Davidson, R. J. (2004). The role of the central nucleus of the amygdala in mediating fear and anxiety in the primate. Journal of Neuroscience, 24, 5506–5515. 10.1523/JNEUROSCI.0292-04.2004 http://ezproxy.library.wisc.edu/login?url=http://search.ebscohost.com/login.aspx?direct=true&db=psyh&AN=2004-15551-005&site=ehost-live

Katzelnick, D. J., Kobak, K. A., DeLeire, T., Henk, H. J., Greist, J. H., Davidson, J. R., … Helstad, C.P.. (2001). Impact of generalized social anxiety disorder in managed care. Am J Psychiatry, 158, 1999–2007. 10.1176/appi.ajp.158.12.1999

Kessler, R. C. (2003). The impairments caused by social phobia in the general population: implications for intervention. Acta Psychiatr Scand, 108 (Suppl. 417), 19–27.

Kessler, R. C., Petukhova, M., Sampson, N. A., Zaslavsky, A. M., & Wittchen, H. U. (2012). Twelve-month and lifetime prevalence and lifetime morbid risk of anxiety and mood disorders in the United States. Int J Methods Psychiatr Res, 21, 169–184. 10.1002/mpr.1359

Kim, H. C., Kaplan, C. M., Islam, S., Anderson, A. S., Piper, M. E., Bradford, D. E., … Shackman, A.J.. (2023). Acute nicotine abstinence amplifies subjective withdrawal symptoms and threat-evoked fear and anxiety, but not extended amygdala reactivity. PLoS One, 18, e0288544. 10.1371/journal.pone.0288544

Kivity, Y., & Huppert, J. D. (2015). Emotional reactions to facial expressions in social anxiety: A meta-analysis of self-reports. Emotion Review, 8, 367–375. 10.1177/1754073915594436

Klumpp, H., & Fitzgerald, J. M. (2018). Neuroimaging predictors and mechanisms of treatment response in Social Anxiety Disorder: An overview of the amygdala. Curr Psychiatry Rep, 20, 89. 10.1007/s11920-018-0948-1

Korn, C. W., Vunder, J., Miró, J., Fuentemilla, L., Hurlemann, R., & Bach, D. R. (2017). Amygdala lesions reduce anxiety-like behavior in a human benzodiazepine-sensitive approach-avoidance conflict test. Biological Psychiatry, 82, 522–531.

Koyuncu, A., İnce, E., Ertekin, E., & Tükel, R. (2019). Comorbidity in social anxiety disorder: diagnostic and therapeutic challenges. Drugs Context, 8, 212573. 10.7573/dic.212573

Kruger, O., Shiozawa, T., Kreifelts, B., Scheffler, K., & Ethofer, T. (2015). Three distinct fiber pathways of the bed nucleus of the stria terminalis to the amygdala and prefrontal cortex. Cortex, 66, 60–68. 10.1016/j.cortex.2015.02.007

Lahey, B. B., Tiemeier, H., & Krueger, R. F. (2022). Seven reasons why binary diagnostic categories should be replaced with empirically sounder and less stigmatizing dimensions [10.1002/jcv2.12108]. JCPP Advances, 2, e12108. 10.1002/jcv2.12108

Lange, M. D., Daldrup, T., Remmers, F., Szkudlarek, H. J., Lesting, J., Guggenhuber, S., … Pape, H.C.. (2017). Cannabinoid CB1 receptors in distinct circuits of the extended amygdala determine fear responsiveness to unpredictable threat. Mol Psychiatry, 22, 1422–1430. 10.1038/mp.2016.156

Lee, Y., Fitz, S., Johnson, P. L., & Shekhar, A. (2008). Repeated stimulation of CRF receptors in the BNST of rats selectively induces social but not panic-like anxiety. Neuropsychopharmacology, 33, 2586–2594. 10.1038/sj.npp.1301674

Lipsitz, J. D., & Schneier, F. R. (2000). Social phobia. Epidemiology and cost of illness. PharmacoEconomics, 18, 23–32. https://www.ncbi.nlm.nih.gov/pubmed/11010601

Lorberbaum, J. P., Kose, S., Johnson, M. R., Arana, G. W., Sullivan, L. K., Hamner, M. B., … George, M.S.. (2004). Neural correlates of speech anticipatory anxiety in generalized social phobia. Neuroreport, 15, 2701–2705.

Lungwitz, E. A., Molosh, A., Johnson, P. L., Harvey, B. P., Dirks, R. C., Dietrich, A., … Truitt, W.A.. (2012). Orexin-A induces anxiety-like behavior through interactions with glutamatergic receptors in the bed nucleus of the stria terminalis of rats. Physiol Behav, 107, 726–732. 10.1016/j.physbeh.2012.05.019

Ma, D. S., Correll, J., & Wittenbrink, B. (2015). The Chicago face database: A free stimulus set of faces and norming data. Behav Res Methods, 47(4), 1122–1135. 10.3758/s13428-014-0532-5

Makol, B. A., Youngstrom, E. A., Racz, S. J., Qasmieh, N., Glenn, L. E., & De Los Reyes, A. (2020). Integrating multiple informants’ reports: How conceptual and measurement models may address long-standing problems in clinical decision-making. Clinical Psychological Science, 8, 953–970. 10.1177/2167702620924439

Mathew, A. R., Pettit, J. W., Lewinsohn, P. M., Seeley, J. R., & Roberts, R. E. (2011). Co-morbidity between major depressive disorder and anxiety disorders: shared etiology or direct causation? Psychol Med, 41, 2023–2034. 10.1017/S0033291711000407

McCormick, M., Liu, X., Jomier, J., Marion, C., & Ibanez, L. (2014). ITK: enabling reproducible research and open science. Frontiers in Neuroinformatics, 8, 13. 10.3389/fninf.2014.00013

Merikangas, K. R., Avenevoli, S., Acharyya, S., Zhang, H., & Angst, J. (2002). The spectrum of social phobia in the Zurich cohort study of young adults. Biological Psychiatry, 51, 81–91. 10.1016/S0006-3223(01)01309-9

Meyer, C., Padmala, S., & Pessoa, L. (2019). Dynamic threat processing. J Cogn Neurosci, 31, 522–542. 10.1162/jocn_a_01363

Michalska, K. J., Benson, B., Ivie, E. J., Sachs, J. F., Haller, S. P., Abend, R., … Pine, D.S.. (2023). Neural responding during uncertain threat anticipation in pediatric anxiety. Int J Psychophysiol, 183, 159–170. 10.1016/j.ijpsycho.2022.07.006

Moscarello, J. M., & Penzo, M. A. (2022). The central nucleus of the amygdala and the construction of defensive modes across the threat-imminence continuum. Nat Neurosci, 25, 999–1008. 10.1038/s41593-022-01130-5

Mumford, J. A., Poline, J. B., & Poldrack, R. A. (2015). Orthogonalization of regressors in FMRI models. PLoS One, 10, e0126255. 10.1371/journal.pone.0126255

Murty, D. V. P. S., Song, S., Surampudi, S. G., & Pessoa, L. (2023). Threat and reward imminence processing in the human brain. J Neurosci, 43, 2973–2987. 10.1523/jneurosci.1778-22.2023

NIMH. (2011). Negative valence systems: Workshop proceedings (March 13, 2011 – March 15, 2011; Rockville, Maryland). Retrieved July 1 from https://www.nimh.nih.gov/research/research-funded-by-nimh/rdoc/negative-valence-systems-workshop-proceedings.shtml

Oler, J. A., Fox, A. S., Shackman, A. J., & Kalin, N. H. (2016). The central nucleus of the amygdala is a critical substrate for individual differences in anxiety. In D. G. Amaral & R. Adolphs (Eds.), Living without an amygdala (pp. 218–251). Guilford.

Petersen, A. C., Crockett, L., Richards, M., & Boxer, A. (1988). (1988). Pubertal Development Scale (PDS). APA PsycTests. 10.1037/t06349-000

Poldrack, R. A., Baker, C. I., Durnez, J., Gorgolewski, K. J., Matthews, P. M., Munafo, M. R., … Yarkoni, T. (2017). Scanning the horizon: towards transparent and reproducible neuroimaging research. Nat Rev Neurosci, 18, 115–126. 10.1038/nrn.2016.167

Poline, J.-B., Kherif, F., & Penny, W. (2007). Contrasts and Classical Inference. Statistical Parametric Mapping: The Analysis of Functional Brain Images. 10.1016/B978-012372560-8/50009-7

Pomrenze, M. B., Giovanetti, S. M., Maiya, R., Gordon, A. G., Kreeger, L. J., & Messing, R. O. (2019). Dissecting the roles of GABA and neuropeptides from rat central amygdala CRF neurons in anxiety and fear learning. Cell Rep, 29, 13–21 e14. 10.1016/j.celrep.2019.08.083

Pomrenze, M. B., Tovar-Diaz, J., Blasio, A., Maiya, R., Giovanetti, S. M., Lei, K., … Messing, R.O.. (2019). A corticotropin releasing factor network in the extended amygdala for anxiety. J Neurosci, 39, 1030–1043. 10.1523/JNEUROSCI.2143-18.2018

Pruim, R. H. R., Mennes, M., van Rooij, D., Llera, A., Buitelaar, J. K., & Beckmann, C. F. (2015). ICA-AROMA: a robust ICA-based strategy for removing motion artifacts from fMRI data. Neuroimage, 112, 267–277.

R Core Team. (2022). R: A language and environment for statistical computing. In. Vienna, Austria: R Foundation for Statistical Computing.

Rapee, R. M., McLellan, L. F., Carl, T., Trompeter, N., Hudson, J. L., Jones, M. P., & Wuthrich, V. M. (2023). Comparison of transdiagnostic treatment and specialized social anxiety treatment for children and adolescents with Social Anxiety Disorder: A randomized controlled trial. J Am Acad Child Adolesc Psychiatry, 62, 646–655. 10.1016/j.jaac.2022.08.003

Ren, J., Lu, C. L., Huang, J., Fan, J., Guo, F., Mo, J. W., … Cao, X. (2022). A distinct metabolically defined central nucleus circuit bidirectionally controls anxiety-related behaviors. J Neurosci, 42, 2356–2370. 10.1523/jneurosci.1578-21.2022

Ressler, R. L., Goode, T. D., Evemy, C., & Maren, S. (2020). NMDA receptors in the CeA and BNST differentially regulate fear conditioning to predictable and unpredictable threats. Neurobiology of Learning and Memory, 174, 107281. 10.1016/j.nlm.2020.107281

RStudio Team. (2022). RStudio: Integrated Development for R. In. Boston, MA: RStudio PBC.

Sajdyk, T., Johnson, P., Fitz, S., & Shekhar, A. (2008). Chronic inhibition of GABA synthesis in the bed nucleus of the stria terminalis elicits anxiety-like behavior. J Psychopharmacol, 22, 633–641. 10.1177/0269881107082902

Schneier, F. R., Blanco, C., Antia, S. X., & Liebowitz, M. R. (2002). The social anxiety spectrum. Psychiatr Clin North Am, 25, 757–774. 10.1016/s0193-953x(02)00018-7

Schneier, F. R., Johnson, J., Hornig, C. D., Liebowitz, M. R., & Weissman, M. M. (1992). Social phobia: comorbidity and morbidity in an epidemiologic sample. Archives of General Psychiatry, 49(4), 282–288.

Scholten, W. D., Batelaan, N. M., Penninx, B. W., van Balkom, A. J., Smit, J. H., Schoevers, R. A., & van Oppen, P. (2016). Diagnostic instability of recurrence and the impact on recurrence rates in depressive and anxiety disorders. J Affect Disord, 195, 185–190. 10.1016/j.jad.2016.02.025

Scholten, W. D., Batelaan, N. M., van Balkom, A. J., Wjh Penninx, B., Smit, J. H., & van Oppen, P. (2013). Recurrence of anxiety disorders and its predictors. J Affect Disord, 147, 180–185. 10.1016/j.jad.2012.10.031

Shackman, A. J., & Fox, A. S. (2016). Contributions of the central extended amygdala to fear and anxiety. Journal of Neuroscience, 36, 8050–8063. 10.1523/JNEUROSCI.0982-16.2016

Shackman, A. J., & Fox, A. S. (2018). Getting serious about variation: Lessons for clinical neuroscience. Trends in cognitive sciences, 22, 368–369.

Shackman, A. J., & Fox, A. S. (2021). Two decades of anxiety neuroimaging research: New insights and a look to the future American Journal of Psychiatry, 178, 106–109.

Shackman, A. J., Fox, A. S., Oler, J. A., Shelton, S. E., Davidson, R. J., & Kalin, N. H. (2013). Neural mechanisms underlying heterogeneity in the presentation of anxious temperament. Proceedings of the National Academy of Sciences of the United States of America, 110, 6145–6150. 1214364110 [pii] 10.1073/pnas.1214364110

Shackman, A. J., Grogans, S. E., & Fox, A. S. (in press). Fear, anxiety, and the functional architecture of the human central extended amygdala. Nature Reviews Neuroscience.

Shackman, A. J., Stockbridge, M. D., Tillman, R. M., Kaplan, C. M., Tromp, D. P. M., Fox, A. S., & Gamer, M. (2016). The neurobiology of anxiety and attentional biases to threat: Implications for understanding anxiety disorders in adults and youth. Journal of Experimental Psychopathology, 7, 311–342.

Sheehan, D. V., Sheehan, K. H., Shytle, R. D., Janavs, J., Bannon, Y., Rogers, J. E., … Wilkinson, B. (2010). Reliability and validity of the Mini International Neuropsychiatric Interview for Children and Adolescents (MINI-KID). J Clin Psychiatry, 71(3), 313–326. 10.4088/JCP.09m05305whi

Singewald, N., Sartori, S. B., Reif, A., & Holmes, A. (2023). Alleviating anxiety and taming trauma: Novel pharmacotherapeutics for anxiety disorders and posttraumatic stress disorder. Neuropharmacology, 226, 109418. 10.1016/j.neuropharm.2023.109418

Sladky, R., Geissberger, N., Pfabigan, D. M., Kraus, C., Tik, M., Woletz, M., … Windischberger, C. (2018). Unsmoothed functional MRI of the human amygdala and bed nucleus of the stria terminalis during processing of emotional faces. Neuroimage, 168, 383–391. 10.1016/j.neuroimage.2016.12.024

Spinhoven, P., Batelaan, N., Rhebergen, D., van Balkom, A., Schoevers, R., & Penninx, B. W. (2016). Prediction of 6-yr symptom course trajectories of anxiety disorders by diagnostic, clinical and psychological variables. J Anxiety Disord, 44, 92–101. 10.1016/j.janxdis.2016.10.011

Stein, D. J., Lim, C. C. W., Roest, A. M., de Jonge, P., Aguilar-Gaxiola, S., Al-Hamzawi, A., … WHO World Mental Health Survey Collaborators. (2017). The cross-national epidemiology of social anxiety disorder: Data from the World Mental Health Survey Initiative. BMC Medicine, 15, 143. 10.1186/s12916-017-0889-2

Straube, T., Mentzel, H. J., & Miltner, W. H. (2005). Common and distinct brain activation to threat and safety signals in social phobia. Neuropsychobiology, 52(3), 163–168. 10.1159/000087987

Strawn, J. R., Lu, L., Peris, T. S., Levine, A., & Walkup, J. T. (2021). Pediatric anxiety disorders – what have we learnt in the last 10 years? Journal of Child Psychology and Psychiatry, 62, 114–139. 10.1111/jcpp.13262

Tan, E., Zeytinoglu, S., Morales, S., Buzzell, G. A., Almas, A. N., Degnan, K. A., … Fox, N.A.. (*in press*). Social versus non-social behavioral inhibition: Differential prediction from early childhood of long-term psychosocial outcomes. Developmental Science.

Theiss, J. D., Ridgewell, C., McHugo, M., Heckers, S., & Blackford, J. U. (2017). Manual segmentation of the human bed nucleus of the stria terminalis using 3T MRI. Neuroimage, 146, 288–292. 10.1016/j.neuroimage.2016.11.047

Tiego, J., Martin, E., DeYoung, C. G., Hagan, K., Cooper, S. E., Pasion, R., … The HiTOP Neurobiological Foundations Work Group. (2023). Precision behavioral phenotyping as a strategy for uncovering the biological correlates of psychopathology. Nature Mental Health, 1, 304–315. doi:10.31219/osf.io/geh6q

Tillfors, M., Furmark, T., Marteinsdottir, I., & Fredrikson, M. (2002). Cerebral blood flow during anticipation of public speaking in social phobia: a PET study. Biol Psychiatry, 52, 1113–1119. 10.1016/s0006-3223(02)01396-3

Tillman, R. M., Stockbridge, M. D., Nacewicz, B. M., Torrisi, S., Fox, A. S., Smith, J. F., & Shackman, A. J. (2018). Intrinsic functional connectivity of the central extended amygdala. Human Brain Mapping, 39, 1291–1312.

Tranel, D., Gullickson, G., Koch, M., & Adolphs, R. (2006). Altered experience of emotion following bilateral amygdala damage. Cognitive Neuropsychiatry, 11, 219–232.

Tseng, Y.-T., Schaefke, B., Wei, P., & Wang, L. (2023). Defensive responses: behaviour, the brain and the body. Nature Reviews Neuroscience, 24, 655–671. 10.1038/s41583-023-00736-3

Tukey, J. W. (1977). Exploratory data analysis. Addison Wesley.

Tulbure, B. T., Szentagotai, A., Dobrean, A., & David, D. (2012). Evidence based clinical assessment of child and adolescent social phobia: a critical review of rating scales. Child Psychiatry Hum Dev, 43, 795–820. 10.1007/s10578-012-0297-y

Tustison, N. J., Avants, B. B., Cook, P. A., Zheng, Y. J., Egan, A., Yushkevich, P. A., & Gee, J. C. (2010). N4ITK: Improved N3 bias correction [Article]. IEEE Transactions on Medical Imaging, 29, 1310–1320. 10.1109/tmi.2010.2046908

Watson, D., Levin-Aspenson, H. F., Waszczuk, M. A., Conway, C. C., Dalgleish, T., Dretsch, M. N., … HiTOP Utility Workgroup. (2022). Validity and utility of Hierarchical Taxonomy of Psychopathology (HiTOP): III. Emotional dysfunction superspectrum. World Psychiatry, 21, 26–54. 10.1002/wps.20943

Weeks, J. W., Howell, A. N., Srivastav, A., & Goldin, P. R. (2019). “Fear guides the eyes of the beholder”: Assessing gaze avoidance in social anxiety disorder via covert eye tracking of dynamic social stimuli. J Anxiety Disord, 65, 56–63. 10.1016/j.janxdis.2019.05.005

Wellcome Centre for Human Neuroimaging. (2022). SPM. University College London. Retrieved April 18 from https://fil.ion.ucl.ac.uk/spm/

Wickham, H. (2016). ggplot2: Elegant graphics for data analysis (2nd ed.). Springer-Verlag.

Wong, Q. J., Gregory, B., & McLellan, L. F. (2016). A review of scales to measure social anxiety disorder in clinical and epidemiological studies. Curr Psychiatry Rep, 18, 38. 10.1007/s11920-016-0677-2

Zhu, Y., Xie, S. Z., Peng, A. B., Yu, X. D., Li, C. Y., Fu, J. Y., … Li, X.M.. (2024). Distinct circuits from the central lateral amygdala to the ventral part of the bed nucleus of stria terminalis regulate different fear memory. Biol Psychiatry, 95, 732–744. 10.1016/j.biopsych.2023.08.022

Ziv, M., Goldin, P. R., Jazaieri, H., Hahn, K. S., & Gross, J. J. (2013). Is there less to social anxiety than meets the eye? Behavioral and neural responses to three socio-emotional tasks. Biol Mood Anxiety Disord, 3, 5. 10.1186/2045-5380-3-5

