## Supplementary Results for "Adolescent social anxiety is associated with diminished discrimination of anticipated threat and safety in the bed nucleus of the stria terminalis"

**Table S1**. Descriptive statistics for clusters and local extrema showing omnibus differences in activation between the three Anticipation conditions (Uncertain Threat, Certain Threat, and Uncertain Safety) relative to implicit baseline (Certain Safety Anticipation), *p*<.05, whole-brain corrected.

| **Cluster** | **Hemisphere** | **Label** | **mm$^3$** | ***F*** | ***x*** | ***y*** | ***z*** |
|---|---|---|---|---|---|---|---|
| 1 | L | Cuneal Cortex | 86,824 | 8.96 | -12 | -80 | 30 |
| 1 | L | Intracalcarine Cortex | 86,824 | 89.58 | -14 | -74 | 8 |
| 1 | L | Lateral Occipital Cortex, superior division | 86,824 | 10.68 | -12 | -84 | 46 |
| 1 | L | Lingual Gyrus | 86,824 | 17.71 | -18 | -52 | -10 |
| 1 | L | Occipital Fusiform Gyrus | 86,824 | 32.06 | -28 | -72 | -12 |
| 1 | L | Occipital Pole | 86,824 | 61.57 | -8 | -98 | -4 |
| 1 | L | Temporal Occipital Fusiform Cortex | 86,824 | 23.56 | -32 | -58 | -14 |
| 1 | R | Cuneal Cortex | 86,824 | 26.25 | 4 | -86 | 34 |
| 1 | R | Intracalcarine Cortex | 86,824 | 130.35 | 14 | -78 | 12 |
| 1 | R | Lingual Gyrus | 86,824 | 17.54 | 18 | -44 | -10 |
| 1 | R | Occipital Fusiform Gyrus | 86,824 | 43.76 | 26 | -76 | -12 |
| 1 | R | Occipital Pole | 86,824 | 69.28 | 8 | -90 | -2 |
| 1 | R | Supracalcarine Cortex | 86,824 | 104.29 | 2 | -80 | 8 |
| 1 | R | Temporal Occipital Fusiform Cortex | 86,824 | 14.71 | 28 | -48 | -12 |
| 2 | L | Postcentral Gyrus | 48,576 | 8.68 | -14 | -46 | 70 |
| 2 | L | Precuneus Cortex | 48,576 | 16.41 | -10 | -50 | 58 |
| 2 | L | Superior Parietal Lobule | 48,576 | 17.66 | -28 | -46 | 60 |
| 2 | R | Angular Gyrus | 48,576 | 17.94 | 60 | -46 | 22 |
| 2 | R | Lateral Occipital Cortex, inferior division | 48,576 | 46.47 | 48 | -72 | 0 |
| 2 | R | Lateral Occipital Cortex, superior division | 48,576 | 23.03 | 48 | -68 | 20 |
| 2 | R | Middle Temporal Gyrus, temporooccipital part | 48,576 | 23.57 | 60 | -46 | 2 |
| 2 | R | Postcentral Gyrus | 48,576 | 14.62 | 42 | -36 | 58 |
| 2 | R | Precuneus Cortex | 48,576 | 16.16 | 8 | -48 | 54 |
| 2 | R | Superior Parietal Lobule | 48,576 | 18.55 | 26 | -50 | 52 |
| 2 | R | Supramarginal Gyrus, posterior division | 48,576 | 17.66 | 44 | -38 | 50 |
| 2 | R | Temporal Occipital Fusiform Cortex | 48,576 | 27.99 | 44 | -50 | -16 |
| 3 | R | Middle Frontal Gyrus | 8,360 | 11.27 | 28 | 12 | 54 |
| 3 | R | Precentral Gyrus | 8,360 | 18.63 | 26 | -12 | 66 |
| 3 | R | Superior Frontal Gyrus | 8,360 | 15.01 | 20 | -2 | 58 |
| 4 | L | Middle Frontal Gyrus | 8,080 | 11.80 | -38 | 4 | 46 |
| 4 | L | Paracingulate Gyrus | 8,080 | 14.49 | -10 | 12 | 44 |
| 4 | L | Precentral Gyrus | 8,080 | 16.97 | -32 | -10 | 56 |
| 4 | L | Superior Frontal Gyrus | 8,080 | 15.40 | -12 | 6 | 64 |
| 4 | R | Midcingulate Cortex | 8,080 | 17.13 | 6 | 8 | 26 |
| 4 | R | Paracingulate Gyrus | 8,080 | 9.35 | 6 | 14 | 40 |
| 5 | R | Inferior Frontal Gyrus, pars opercularis | 7,576 | 21.36 | 40 | 12 | 26 |

| | | | | | | | |
|---|---|---|---|---|---|---|---|
| 5 | R | Precentral Gyrus | 7,576 | 30.19 | 46 | 8 | 32 |
| 6 | R | Frontal Pole/ dorsolateral Prefronal Cortex | 3,216 | 14.79 | 26 | 42 | 28 |
| 6 | R | Middle Frontal Gyrus | 3,216 | 12.03 | 32 | 34 | 32 |
| 7 | L | Inferior Temporal Gyrus, temporooccipital part | 2,800 | 15.91 | -46 | -62 | -12 |
| 7 | L | Middle Temporal Gyrus, posterior division | 2,800 | 10.41 | -58 | -44 | -2 |
| 7 | L | Middle Temporal Gyrus, temporooccipital part | 2,800 | 12.94 | -56 | -50 | 2 |
| 8 | L | Midcingulate Cortex | 1,096 | 11.69 | -10 | 24 | 26 |
| 9 | L | Lateral Occipital Cortex, superior division | 1,096 | 13.78 | -22 | -76 | 36 |

**Table S2.** Descriptive statistics for clusters and local extrema showing greater activation during the uncertain versus certain anticipation of threatening faces and voices (masked by the omnibus *F*-test), *p*<.05, whole-brain corrected.

| Cluster | Hemisphere | Label | mm³ | *t* | *x* | *y* | *z* |
|---|---|---|---|---|---|---|---|
| 1 | R | Angular Gyrus | 38,152 | 5.57 | 62 | -46 | 18 |
| 1 | R | Lateral Occipital Cortex, inferior division | 38,152 | 8.48 | 52 | -72 | -2 |
| 1 | R | Lateral Occipital Cortex, superior division | 38,152 | 4.95 | 22 | -68 | 42 |
| 1 | R | Middle Temporal Gyrus, temporooccipital part | 38,152 | 6.95 | 60 | -46 | 2 |
| 1 | R | Postcentral Gyrus | 38,152 | 4.87 | 42 | -36 | 58 |
| 1 | R | Superior Parietal Lobule | 38,152 | 5.17 | 24 | -50 | 54 |
| 1 | R | Supramarginal Gyrus, posterior division | 38,152 | 5.78 | 42 | -40 | 48 |
| 1 | R | Temporal Occipital Fusiform Cortex | 38,152 | 6.74 | 44 | -50 | -18 |
| 2 | L | Postcentral Gyrus | 8,064 | 4.22 | -14 | -46 | 70 |
| 2 | L | Precuneus Cortex | 8,064 | 5.65 | -10 | -50 | 56 |
| 2 | L | Superior Parietal Lobule | 8,064 | 6.00 | -28 | -46 | 60 |
| 2 | R | Lateral Occipital Cortex, superior division | 8,064 | 4.47 | 14 | -62 | 54 |
| 2 | R | Precuneus Cortex | 8,064 | 5.72 | 10 | -46 | 52 |
| 3 | R | Middle Frontal Gyrus | 7,624 | 4.78 | 28 | 12 | 54 |
| 3 | R | Precentral Gyrus | 7,624 | 5.11 | 38 | -6 | 50 |
| 3 | R | Superior Frontal Gyrus | 7,624 | 5.14 | 22 | 4 | 52 |
| 4 | R | Inferior Frontal Gyrus, pars opercularis | 7,576 | 6.33 | 40 | 12 | 26 |
| 4 | R | Precentral Gyrus | 7,576 | 7.63 | 44 | 6 | 32 |
| 5 | L | Middle Frontal Gyrus | 7,432 | 4.76 | -40 | 4 | 46 |
| 5 | L | Paracingulate Gyrus | 7,432 | 5.46 | -10 | 12 | 44 |
| 5 | L | Precentral Gyrus | 7,432 | 5.38 | -32 | -8 | 56 |
| 5 | L | Superior Frontal Gyrus | 7,432 | 4.81 | -26 | 0 | 62 |
| 5 | R | Midcingulate Cortex | 7,432 | 4.89 | 8 | 24 | 28 |
| 5 | R | Paracingulate Gyrus | 7,432 | 4.08 | 2 | 12 | 42 |
| 6 | R | Frontal Pole | 3,184 | 5.29 | 26 | 42 | 28 |
| 7 | L | Inferior Temporal Gyrus, temporooccipital part | 2,648 | 5.49 | -46 | -62 | -12 |
| 7 | L | Middle Temporal Gyrus, posterior division | 2,648 | 4.32 | -58 | -44 | -2 |
| 7 | L | Middle Temporal Gyrus, temporooccipital part | 2,648 | 4.87 | -56 | -50 | 4 |
| 8 | L | Midcingulate Cortex | 1,080 | 4.79 | -10 | 24 | 26 |
| 9 | L | Lateral Occipital Cortex, superior division | 1,072 | 5.29 | -24 | -76 | 36 |
| 10 | R | Midcingulate Cortex | 384 | 5.29 | 4 | 12 | 24 |
| 11 | L | Cuneal Cortex | 40 | 3.68 | -12 | -82 | 30 |

**Table S3**. Descriptive statistics for clusters and local extrema showing greater activation during the certain versus uncertain anticipation of threatening faces and voices (masked by the omnibus *F*-test), $p<.05$, whole-brain corrected.

| **Cluster** | **Hemisphere** | **Label** | **mm³** | ***t*** | ***x*** | ***y*** | ***z*** |
|---|---|---|---|---|---|---|---|
| 1 | L | Intracalcarine Cortex | 82,288 | 11.82 | -14 | -74 | 6 |
| 1 | L | Lateral Occipital Cortex, inferior division | 82,288 | 4.06 | -38 | -86 | -16 |
| 1 | L | Lingual Gyrus | 82,288 | 5.57 | -18 | -52 | -10 |
| 1 | L | Occipital Fusiform Gyrus | 82,288 | 7.64 | -28 | -76 | -12 |
| 1 | L | Occipital Pole | 82,288 | 11.13 | -8 | -98 | -4 |
| 1 | L | Temporal Occipital Fusiform Cortex | 82,288 | 6.57 | -32 | -58 | -14 |
| 1 | R | Cuneal Cortex | 82,288 | 7.03 | 2 | -86 | 36 |
| 1 | R | Intracalcarine Cortex | 82,288 | 11.50 | 12 | -88 | 2 |
| 1 | R | Occipital Fusiform Gyrus | 82,288 | 9.19 | 26 | -76 | -12 |
| 1 | R | Occipital Pole | 82,288 | 11.94 | 6 | -90 | -2 |
| 1 | R | Supracalcarine Cortex | 82,288 | 12.56 | 2 | -80 | 8 |
| 2 | R | Lingual Gyrus | 504 | 4.98 | 18 | -44 | -10 |
| 2 | R | Temporal Occipital Fusiform Cortex | 504 | 3.33 | 26 | -48 | -14 |

**Table S4**. Descriptive statistics for clusters and local extrema showing greater activation during the uncertain anticipation of threatening versus benign faces and voices (masked by the omnibus *F*-test), *p*<.05, whole-brain corrected.

| **Cluster** | **Hemisphere** | **Label** | **mm$^3$** | ***t*** | ***x*** | ***y*** | ***z*** |
|---|---|---|---|---|---|---|---|
| 1 | L | Precuneus Cortex | 3,640 | 5.16 | -6 | -50 | 54 |
| 1 | R | Cingulate Gyrus, posterior division | 3,640 | 3.82 | 10 | -32 | 40 |
| 1 | R | Precuneus Cortex | 3,640 | 4.91 | 6 | -46 | 54 |
| 2 | R | Angular Gyrus | 880 | 5.03 | 60 | -46 | 22 |
| 3 | L | Lateral Occipital Cortex, superior division | 544 | 4.06 | -26 | -80 | 34 |
| 4 | L | Lateral Occipital Cortex, superior division | 16 | 3.48 | -18 | -88 | 36 |

**Table S5**. Descriptive statistics for clusters and local extrema showing greater activation during the uncertain anticipation of benign versus threatening faces and voices (masked by the omnibus *F*-test), $p$<.05, whole-brain corrected.

| **Cluster** | **Hemisphere** | **Label** | **mm³** | ***t*** | ***x*** | ***y*** | ***z*** |
|---|---|---|---|---|---|---|---|
| 1 | L | Intracalcarine Cortex | 57,272 | 12.50 | -12 | -74 | 10 |
| 1 | L | Occipital Pole | 57,272 | 7.38 | -2 | -94 | -4 |
| 1 | R | Cuneal Cortex | 57,272 | 5.92 | 2 | -84 | 36 |
| 1 | R | Intracalcarine Cortex | 57,272 | 16.37 | 14 | -78 | 12 |
| 1 | R | Occipital Pole | 57,272 | 6.87 | 14 | -102 | -2 |
| 2 | R | Occipital Fusiform Gyrus | 992 | 5.28 | 30 | -68 | -12 |
| 2 | R | Temporal Occipital Fusiform Cortex | 992 | 4.14 | 34 | -60 | -12 |
| 3 | L | Lingual Gyrus | 368 | 5.55 | -16 | -52 | -12 |
| 4 | R | Lateral Occipital Cortex, inferior division | 8 | 3.37 | 40 | -86 | -10 |

**Table S6**. Descriptive statistics for clusters and local extrema showing greater activation during the certain anticipation of threatening faces and voices versus the uncertain anticipation of benign faces and voices (masked by the omnibus *F*-test), $p<.05$, whole-brain corrected.

| **Cluster** | **Hemisphere** | **Label** | **mm³** | ***t*** | ***x*** | ***y*** | ***z*** |
|---|---|---|---|---|---|---|---|
| 1 | L | Lingual Gyrus | 29,288 | 7.55 | -6 | -90 | -10 |
| 1 | L | Occipital Fusiform Gyrus | 29,288 | 6.90 | -28 | -70 | -12 |
| 1 | L | Occipital Pole | 29,288 | 5.25 | -8 | -98 | 2 |
| 1 | L | Temporal Occipital Fusiform Cortex | 29,288 | 6.26 | -32 | -56 | -14 |
| 1 | R | Cuneal Cortex | 29,288 | 3.31 | 6 | -86 | 38 |
| 1 | R | Lingual Gyrus | 29,288 | 7.98 | 6 | -90 | -6 |
| 1 | R | Occipital Fusiform Gyrus | 29,288 | 8.24 | 24 | -76 | -10 |
| 1 | R | Occipital Pole | 29,288 | 6.75 | 14 | -98 | 2 |
| 1 | R | Temporal Occipital Fusiform Cortex | 29,288 | 5.49 | 28 | -48 | -12 |
| 2 | L | Lateral Occipital Cortex, superior division | 1,192 | 3.61 | -12 | -84 | 46 |
| 2 | L | Occipital Pole | 1,192 | 4.94 | -14 | -90 | 34 |
| 3 | L | Precuneus Cortex | 56 | 4.08 | -4 | -82 | 42 |

**Table S7.** Descriptive statistics for clusters and local extrema showing greater activation during the uncertain anticipation of benign faces and voices versus the certain anticipation of threatening faces and voices (masked by the omnibus *F*-test), $p<.05$, whole-brain corrected.

| Cluster | Hemisphere | Label | mm³ | *t* | *x* | *y* | *z* |
|---|---|---|---|---|---|---|---|
| 1 | R | Inferior Temporal Gyrus, temporooccipital part | 21,888 | 4.84 | 56 | -52 | -12 |
| 1 | R | Lateral Occipital Cortex, inferior division | 21,888 | 9.05 | 46 | -74 | 0 |
| 1 | R | Middle Temporal Gyrus, temporooccipital part | 21,888 | 5.74 | 54 | -56 | 0 |
| 1 | R | Temporal Occipital Fusiform Cortex | 21,888 | 6.02 | 46 | -56 | -18 |
| 2 | R | Lateral Occipital Cortex, superior division | 5,040 | 4.00 | 38 | -66 | 34 |
| 2 | R | Postcentral Gyrus | 5,040 | 4.11 | 14 | -44 | 62 |
| 2 | R | Superior Parietal Lobule | 5,040 | 5.14 | 32 | -40 | 64 |
| 2 | R | Supramarginal Gyrus, posterior division | 5,040 | 4.27 | 42 | -40 | 50 |
| 3 | R | Inferior Frontal Gyrus, pars opercularis | 4,136 | 3.35 | 54 | 12 | 16 |
| 4 | R | Intracalcarine Cortex | 2,640 | 5.20 | 14 | -76 | 12 |
| 5 | L | Intracalcarine Cortex | 2,408 | 4.37 | -14 | -76 | 12 |
| 6 | R | Precentral Gyrus | 2,352 | 5.54 | 12 | -16 | 74 |
| 7 | L | Lateral Occipital Cortex, inferior division | 2,000 | 3.94 | -48 | -66 | -18 |
| 7 | L | Middle Temporal Gyrus, posterior division | 2,000 | 3.64 | -56 | -42 | -2 |
| 7 | L | Middle Temporal Gyrus, temporooccipital part | 2,000 | 4.30 | -54 | -50 | 4 |
| 7 | L | Temporal Occipital Fusiform Cortex | 2,000 | 4.20 | -42 | -60 | -12 |
| 8 | L | Precentral Gyrus | 768 | 5.30 | -32 | -12 | 56 |
| 9 | L | Superior Frontal Gyrus | 624 | 5.07 | -10 | 6 | 66 |

**Table S8**. Descriptive statistics for clusters and local extrema showing greater activation during the acute presentation of threatening faces and voices versus baseline, $p<.05$, whole-brain corrected.

| Cluster | Hemisphere | Label | mm³ | *t* | *x* | *y* | *z* |
|---|---|---|---|---|---|---|---|
| 1 | R | Inferior Frontal Gyrus, pars triangularis | 359,264 | 8.23 | 50 | 28 | 0 |
| 1 | L | Frontal Operculum Cortex | 359,264 | 10.89 | -38 | 26 | 2 |
| 1 | L | Frontal Orbital Cortex | 359,264 | 11.11 | -36 | 26 | -4 |
| 1 | L | Inferior Frontal Gyrus, pars opercularis | 359,264 | 7.63 | -48 | 20 | 16 |
| 1 | R | Frontal Orbital Cortex | 359,264 | 7.72 | 32 | 18 | -18 |
| 1 | L | Insular Cortex | 359,264 | 7.73 | -30 | 16 | -14 |
| 1 | R | Inferior Frontal Gyrus, pars opercularis | 359,264 | 9.95 | 42 | 14 | 28 |
| 1 | L | Putamen | 359,264 | 5.82 | -20 | 10 | -6 |
| 1 | R | Putamen | 359,264 | 5.15 | 18 | 10 | -2 |
| 1 | L | Temporal Pole | 359,264 | 15.29 | -54 | 8 | -12 |
| 1 | L | Caudate | 359,264 | 3.81 | -8 | 8 | 6 |
| 1 | R | Temporal Pole | 359,264 | 16.33 | 52 | 8 | -14 |
| 1 | R | Precentral Gyrus | 359,264 | 6 | 50 | 4 | 52 |
| 1 | R | Superior Temporal Gyrus, anterior division | 359,264 | 15.68 | 56 | 4 | -14 |
| 1 | R | Cingulate Gyrus, anterior division | 359,264 | 5.3 | 6 | 2 | 28 |
| 1 | L | Superior Temporal Gyrus, anterior division | 359,264 | 17.11 | -58 | -4 | -6 |
| 1 | R | Amygdala | 359,264 | 7.94 | 30 | -4 | -20 |
| 1 | L | Amygdala | 359,264 | 8.17 | -30 | -6 | -22 |
| 1 | L | Amygdala | 359,264 | 8.22 | -16 | -6 | -14 |
| 1 | R | Amygdala | 359,264 | 8.51 | 16 | -6 | -14 |
| 1 | R | Amygdala | 359,264 | 7.43 | 28 | -8 | -14 |
| 1 | R | Planum Temporale | 359,264 | 17.87 | 62 | -10 | 2 |
| 1 | L | Amygdala | 359,264 | 6.85 | -26 | -12 | -12 |
| 1 | R | Heschls Gyrus (includes H1 and H2) | 359,264 | 15.44 | 50 | -14 | 4 |
| 1 | R | Superior Temporal Gyrus, posterior division | 359,264 | 14.11 | 68 | -18 | 6 |
| 1 | L | Heschls Gyrus (includes H1 and H2) | 359,264 | 18.13 | -48 | -20 | 6 |
| 1 | L | Planum Temporale | 359,264 | 17.04 | -60 | -22 | 4 |
| 1 | L | Inferior Temporal Gyrus, posterior division | 359,264 | 6.33 | -44 | -26 | -20 |
| 1 | R | Brain Stem | 359,264 | 11.88 | 12 | -26 | -10 |
| 1 | L | Thalamus | 359,264 | 12.47 | -18 | -30 | -4 |
| 1 | L | Brain Stem | 359,264 | 9.42 | -8 | -30 | -6 |

| | | | | | | | |
|---|---|---|---|---|---|---|---|
| 1 | R | Thalamus | 359,264 | 10.09 | 10 | -30 | -2 |
| 1 | R | Temporal Fusiform Cortex, posterior division | 359,264 | 12 | 38 | -38 | -24 |
| 1 | L | Supramarginal Gyrus, posterior division | 359,264 | 15.34 | -66 | -42 | 12 |
| 1 | L | Lingual Gyrus | 359,264 | 6.05 | -16 | -46 | -8 |
| 1 | R | Lingual Gyrus | 359,264 | 9.11 | 18 | -46 | -10 |
| 1 | L | Temporal Occipital Fusiform Cortex | 359,264 | 13.73 | -38 | -48 | -20 |
| 1 | R | Temporal Occipital Fusiform Cortex | 359,264 | 16.56 | 36 | -56 | -16 |
| 1 | L | Intracalcarine Cortex | 359,264 | 11.22 | -20 | -68 | 6 |
| 1 | R | Precuneus Cortex | 359,264 | 5.76 | 12 | -74 | 40 |
| 1 | R | Occipital Fusiform Gyrus | 359,264 | 15.57 | 30 | -76 | -10 |
| 1 | R | Cuneal Cortex | 359,264 | 6.57 | 4 | -80 | 36 |
| 1 | L | Occipital Fusiform Gyrus | 359,264 | 16.68 | -26 | -84 | -12 |
| 1 | L | Occipital Pole | 359,264 | 17.4 | -14 | -96 | -12 |
| 1 | R | Occipital Pole | 359,264 | 18.68 | 14 | -98 | 0 |
| 2 | L | Paracingulate Gyrus | 12,936 | 7.09 | -10 | 20 | 38 |
| 2 | L | Superior Frontal Gyrus | 12,936 | 5.25 | -10 | 20 | 68 |
| 2 | R | Paracingulate Gyrus | 12,936 | 4.26 | 10 | 20 | 40 |
| 2 | R | Superior Frontal Gyrus | 12,936 | 5.14 | 8 | 18 | 68 |
| 3 | R | Frontal Pole | 2,784 | 3.99 | 10 | 60 | 36 |
| 3 | L | Superior Frontal Gyrus | 2,784 | 6.1 | -2 | 56 | 32 |
| 3 | R | Superior Frontal Gyrus | 2,784 | 4.83 | 6 | 50 | 40 |
| 4 | L | Frontal Pole | 2,008 | 3.77 | -4 | 58 | -18 |
| 4 | B | Frontal Medial Cortex | 2,008 | 5.12 | 0 | 42 | -24 |
| 4 | L | Frontal Medial Cortex | 2,008 | 5.31 | -2 | 32 | -28 |

**Table S9**. Descriptive statistics for clusters and local extrema showing greater activation during the acute presentation of benign faces and voices versus baseline, *p*<.05, whole-brain corrected.

| Cluster | Hemisphere | Label | $mm^3$ | *t* | *x* | *y* | *z* |
|---|---|---|---|---|---|---|---|
| 1 | L | Frontal Orbital Cortex | 313,816 | 9.69 | -34 | 28 | 0 |
| 1 | L | Insular Cortex | 313,816 | 9.61 | -30 | 26 | 0 |
| 1 | R | Inferior Frontal Gyrus, pars triangularis | 313,816 | 9.00 | 52 | 26 | 20 |
| 1 | L | Inferior Frontal Gyrus, pars triangularis | 313,816 | 6.29 | -52 | 24 | 4 |
| 1 | R | Frontal Orbital Cortex | 313,816 | 7.08 | 42 | 24 | -8 |
| 1 | L | Inferior Frontal Gyrus, pars opercularis | 313,816 | 7.95 | -50 | 18 | 24 |
| 1 | R | Inferior Frontal Gyrus, pars opercularis | 313,816 | 10.43 | 48 | 18 | 28 |
| 1 | R | Middle Frontal Gyrus | 313,816 | 10.37 | 46 | 16 | 30 |
| 1 | R | Temporal Pole | 313,816 | 14.54 | 48 | 14 | -18 |
| 1 | L | Temporal Pole | 313,816 | 18.12 | -54 | 6 | -10 |
| 1 | R | Precentral Gyrus | 313,816 | 4.83 | 46 | 4 | 46 |
| 1 | R | Superior Temporal Gyrus, anterior division | 313,816 | 18.13 | 56 | 4 | -14 |
| 1 | R | Amygdala | 313,816 | 7.46 | 26 | 2 | -18 |
| 1 | L | Amygdala | 313,816 | 6.95 | -28 | 0 | -20 |
| 1 | R | Amygdala | 313,816 | 7.43 | 30 | -2 | -24 |
| 1 | R | Amygdala | 313,816 | 8.00 | 32 | -4 | -20 |
| 1 | L | Superior Temporal Gyrus, anterior division | 313,816 | 17.79 | -56 | -6 | -4 |
| 1 | L | Amygdala | 313,816 | 9.18 | -16 | -6 | -14 |
| 1 | L | Amygdala | 313,816 | 7.58 | -30 | -6 | -22 |
| 1 | R | Amygdala | 313,816 | 5.37 | 28 | -12 | -12 |
| 1 | R | Superior Temporal Gyrus, posterior division | 313,816 | 15.69 | 50 | -16 | -6 |
| 1 | R | Heschls Gyrus (includes H1 and H2) | 313,816 | 13.72 | 52 | -18 | 6 |
| 1 | L | Heschls Gyrus (includes H1 and H2) | 313,816 | 15.94 | -46 | -20 | 6 |
| 1 | L | Planum Temporale | 313,816 | 16.62 | -60 | -22 | 6 |
| 1 | R | Brain-Stem | 313,816 | 10.19 | 10 | -26 | -8 |
| 1 | L | Thalamus | 313,816 | 12.39 | -20 | -30 | -4 |
| 1 | R | Thalamus | 313,816 | 9.41 | 10 | -30 | -2 |
| 1 | L | Parietal Operculum Cortex | 313,816 | 4.79 | -34 | -34 | 22 |
| 1 | L | Cingulate Gyrus, posterior division | 313,816 | 5.25 | -6 | -38 | 26 |
| 1 | R | Temporal Fusiform Cortex, posterior division | 313,816 | 11.52 | 38 | -38 | -24 |
| 1 | R | Supramarginal Gyrus, posterior division | 313,816 | 13.61 | 54 | -40 | 10 |

| | | | | | | | |
|---|---|---|---|---|---|---|---|
| 1 | L | Supramarginal Gyrus, posterior division | 313,816 | 12.37 | -66 | -42 | 12 |
| 1 | L | Temporal Fusiform Cortex, posterior division | 313,816 | 12.04 | -40 | -42 | -22 |
| 1 | R | Lingual Gyrus | 313,816 | 6.34 | 18 | -48 | -8 |
| 1 | L | Temporal Occipital Fusiform Cortex | 313,816 | 12.77 | -36 | -50 | -18 |
| 1 | L | Lingual Gyrus | 313,816 | 4.72 | -16 | -50 | -8 |
| 1 | R | Temporal Occipital Fusiform Cortex | 313,816 | 14.47 | 36 | -54 | -16 |
| 1 | R | Precuneus Cortex | 313,816 | 4.97 | 14 | -66 | 40 |
| 1 | R | Intracalcarine Cortex | 313,816 | 6.19 | 12 | -68 | 14 |
| 1 | L | Intracalcarine Cortex | 313,816 | 9.99 | -20 | -70 | 8 |
| 1 | R | Occipital Fusiform Gyrus | 313,816 | 16.57 | 32 | -78 | -12 |
| 1 | R | Cuneal Cortex | 313,816 | 4.36 | 2 | -80 | 36 |
| 1 | L | Occipital Fusiform Gyrus | 313,816 | 15.49 | -26 | -84 | -12 |
| 1 | L | Occipital Pole | 313,816 | 16.92 | -10 | -94 | -10 |
| 1 | R | Occipital Pole | 313,816 | 17.94 | 14 | -98 | 0 |
| 2 | R | Superior Frontal Gyrus | 10,680 | 5.69 | 6 | 24 | 58 |
| 2 | R | Paracingulate Gyrus | 10,680 | 3.57 | 10 | 24 | 32 |
| 2 | L | Paracingulate Gyrus | 10,680 | 5.18 | -10 | 20 | 38 |
| 2 | L | Superior Frontal Gyrus | 10,680 | 4.01 | -10 | 18 | 68 |
| 3 | R | Thalamus | 968 | 6.33 | 6 | -8 | 6 |

**Table S10**. Descriptive statistics for clusters and local extrema showing greater activation during the acute presentation of threatening versus benign faces and voices, *p*<.05, whole-brain corrected.

| Cluster | Hemisphere | Label | $mm^3$ | *t* | *x* | *y* | *z* |
|---|---|---|---|---|---|---|---|
| 1 | L | Intracalcarine Cortex | 54,808 | 7.60 | -16 | -70 | 6 |
| 1 | L | Lateral Occipital Cortex, inferior division | 54,808 | 5.06 | -40 | -72 | -8 |
| 1 | L | Occipital Fusiform Gyrus | 54,808 | 5.21 | -34 | -68 | -18 |
| 1 | L | Occipital Pole | 54,808 | 5.96 | -4 | -100 | -2 |
| 1 | L | Temporal Occipital Fusiform Cortex | 54,808 | 4.88 | -32 | -56 | -16 |
| 1 | R | Cuneal Cortex | 54,808 | 4.56 | 18 | -74 | 30 |
| 1 | R | Intracalcarine Cortex | 54,808 | 6.73 | 14 | -66 | 10 |
| 1 | R | Lateral Occipital Cortex, inferior division | 54,808 | 4.75 | 36 | -86 | -10 |
| 1 | R | Lingual Gyrus | 54,808 | 6.55 | 4 | -88 | -2 |
| 1 | R | Occipital Fusiform Gyrus | 54,808 | 6.23 | 36 | -68 | -14 |
| 1 | R | Occipital Pole | 54,808 | 7.00 | 4 | -90 | 4 |
| 1 | R | Precuneus Cortex | 54,808 | 5.64 | 18 | -72 | 38 |
| 1 | R | Temporal Fusiform Cortex, posterior division | 54,808 | 4.05 | 36 | -36 | -24 |
| 1 | R | Temporal Occipital Fusiform Cortex | 54,808 | 5.82 | 36 | -54 | -14 |
| 2 | R | Planum Temporale | 10,504 | 5.35 | 38 | -32 | 14 |
| 2 | R | Superior Temporal Gyrus, posterior division | 10,504 | 6.50 | 50 | -16 | -6 |
| 2 | R | Supramarginal Gyrus, posterior division | 10,504 | 4.75 | 68 | -38 | 16 |
| 3 | L | Central Opercular Cortex | 7,128 | 3.51 | -62 | -14 | 10 |
| 3 | L | Heschls Gyrus (includes H1 and H2) | 7,128 | 4.75 | -44 | -24 | 8 |
| 3 | L | Planum Temporale | 7,128 | 5.77 | -38 | -34 | 10 |
| 3 | L | Supramarginal Gyrus, posterior division | 7,128 | 4.28 | -66 | -42 | 18 |
| 4 | L | Frontal Orbital Cortex | 3,264 | 5.07 | -28 | 14 | -18 |
| 4 | L | Left Putamen | 3,264 | 4.81 | -22 | 8 | -8 |
| 5 | L | Cingulate Gyrus, posterior division | 2,832 | 5.51 | -2 | -16 | 40 |
| 5 | R | Cingulate Gyrus, posterior division | 2,832 | 5.49 | 6 | -24 | 28 |
| 6 | R | Temporal Pole | 2,816 | 5.44 | 54 | 8 | -16 |
| 7 | L | Brain-Stem | 2,256 | 5.05 | -6 | -36 | -6 |
| 7 | R | Brain-Stem | 2,256 | 4.51 | 12 | -26 | -10 |
| 8 | L | Left Caudate | 1,992 | 3.60 | -8 | 8 | 14 |
| 9 | L | Frontal Pole | 1,416 | 4.24 | -6 | 58 | 18 |
| 10 | R | Frontal Orbital Cortex | 1,400 | 4.93 | 32 | 18 | -20 |
| 10 | R | Right Putamen | 1,400 | 4.47 | 26 | 12 | -6 |
| 11 | R | Precentral Gyrus | 1,200 | 5.15 | 52 | -2 | 48 |
| 12 | L | Inferior Frontal Gyrus, pars opercularis | 800 | 3.98 | -48 | 20 | 10 |

**Table S11**. Descriptive statistics for clusters and local extrema showing greater activation during the acute presentation of benign versus threatening faces and voices, $p$<.05, whole-brain corrected.

| Cluster | Hemisphere | Label | mm$^3$ | *t* | *x* | *y* | *z* |
|---|---|---|---|---|---|---|---|
| 1 | L | Angular Gyrus | 8,464 | 3.85 | -44 | -60 | 24 |
| 1 | L | Lateral Occipital Cortex, superior division | 8,464 | 6.42 | -42 | -78 | 30 |
| 2 | L | Middle Frontal Gyrus | 6,680 | 3.94 | -38 | 10 | 62 |
| 2 | L | Superior Frontal Gyrus | 6,680 | 5.44 | -24 | 14 | 60 |
| 3 | L | Precuneus Cortex | 5,968 | 6.86 | -6 | -54 | 16 |
| 3 | R | Precuneus Cortex | 5,968 | 4.82 | 8 | -54 | 6 |
| 4 | L | Cingulate Gyrus, posterior division | 4,576 | 5.38 | -6 | -38 | 36 |
| 4 | L | Precuneus Cortex | 4,576 | 4.74 | -2 | -54 | 48 |
| 4 | R | Lateral Occipital Cortex, superior division | 4,576 | 3.84 | 12 | -60 | 60 |
| 4 | R | Precuneus Cortex | 4,576 | 5.40 | 10 | -60 | 50 |
| 4 | R | Superior Parietal Lobule | 4,576 | 3.80 | 12 | -54 | 66 |
| 5 | R | Lateral Occipital Cortex, superior division | 2,920 | 6.61 | 46 | -70 | 34 |
| 6 | L | Lateral Occipital Cortex, superior division | 2,896 | 4.55 | -8 | -64 | 62 |
| 6 | L | Postcentral Gyrus | 2,896 | 3.99 | -8 | -48 | 70 |
| 6 | L | Precuneus Cortex | 2,896 | 4.23 | -8 | -66 | 56 |
| 7 | L | Middle Temporal Gyrus, temporooccipital part | 2,208 | 5.62 | -58 | -50 | -6 |
| 8 | L | Parahippocampal Gyrus, posterior division | 2,128 | 5.20 | -28 | -30 | -18 |
| 8 | L | Temporal Fusiform Cortex, posterior division | 2,128 | 5.83 | -26 | -38 | -18 |
| 9 | R | Frontal Pole | 1,232 | 4.12 | 28 | 36 | 50 |
| 9 | R | Superior Frontal Gyrus | 1,232 | 4.11 | 22 | 30 | 42 |
| 10 | R | Middle Frontal Gyrus | 1,000 | 3.96 | 26 | 16 | 54 |

**Table S12.** Descriptive statistics for clusters and local extrema showing greater activation during the temporally uncertain versus certain presentation of benign faces and voices, *p*<.05, whole-brain corrected.

| Cluster | Hemisphere | Label | mm³ | *t* | *x* | *y* | *z* |
|---|---|---|---|---|---|---|---|
| 1 | L | Inferior Temporal Gyrus, temporooccipital part | 6,664 | 5.96 | -42 | -54 | -8 |
| 1 | L | Lateral Occipital Cortex, inferior division | 6,664 | 4.84 | -40 | -70 | -6 |
| 1 | L | Temporal Occipital Fusiform Cortex | 6,664 | 5.78 | -34 | -48 | -16 |
| 2 | L | Superior Frontal Gyrus | 4,200 | 4.29 | -4 | 36 | 46 |
| 2 | R | Cingulate Gyrus, anterior division | 4,200 | 3.60 | 6 | 16 | 34 |
| 2 | B | Juxtapositional Lobule Cortex | 4,200 | 3.93 | 0 | 6 | 48 |
| 2 | R | Paracingulate Gyrus | 4,200 | 4.90 | 8 | 18 | 44 |
| 3 | L | Occipital Pole | 4,176 | 3.69 | -2 | -90 | 26 |
| 3 | L | Precuneus Cortex | 4,176 | 4.23 | -10 | -68 | 42 |
| 3 | R | Lateral Occipital Cortex, superior division | 4,176 | 3.52 | 10 | -84 | 44 |
| 3 | R | Occipital Pole | 4,176 | 4.80 | 18 | -98 | 24 |
| 3 | R | Precuneus Cortex | 4,176 | 3.62 | 4 | -76 | 48 |
| 4 | R | Superior Parietal Lobule | 3,424 | 4.58 | 30 | -42 | 42 |
| 4 | R | Supramarginal Gyrus, posterior division | 3,424 | 4.98 | 48 | -38 | 48 |
| 5 | R | Lateral Occipital Cortex, inferior division | 2,696 | 3.37 | 40 | -78 | -12 |
| 5 | R | Temporal Occipital Fusiform Cortex | 2,696 | 5.23 | 36 | -48 | -18 |
| 6 | R | Precuneus Cortex | 2,504 | 5.06 | 10 | -72 | 36 |
| 7 | L | Precentral Gyrus | 2,328 | 4.54 | -48 | 0 | 36 |
| 8 | R | Frontal Operculum Cortex | 1,424 | 4.61 | 48 | 12 | 0 |
| 8 | R | Insular Cortex | 1,424 | 4.20 | 34 | 20 | 6 |
| 9 | L | Frontal Operculum Cortex | 1,344 | 3.83 | -40 | 14 | 2 |
| 9 | L | Insular Cortex | 1,344 | 4.81 | -30 | 18 | 10 |
| 10 | R | Precentral Gyrus | 1,176 | 4.55 | 48 | 6 | 32 |

**Table S13.** Descriptive statistics for clusters and local extrema showing greater activation during the temporally certain versus uncertain presentation of benign faces and voices, *p*<.05, whole-brain corrected.

| **Cluster** | **Hemisphere** | **Label** | **mm$^3$** | ***t*** | ***x*** | ***y*** | ***z*** |
|---|---|---|---|---|---|---|---|
| 1 | R | Planum Temporale | 2,640 | 4.60 | 58 | -16 | 4 |
| 1 | R | Superior Temporal Gyrus, anterior division | 2,640 | 4.17 | 62 | 0 | -6 |
| 1 | R | Superior Temporal Gyrus, posterior division | 2,640 | 4.61 | 68 | -22 | 4 |
| 2 | L | Heschls Gyrus (includes H1 and H2) | 2,008 | 4.81 | -52 | -18 | 4 |
| 2 | L | Planum Temporale | 2,008 | 3.90 | -62 | -18 | 4 |
| 2 | L | Superior Temporal Gyrus, anterior division | 2,008 | 4.43 | -62 | -6 | -6 |
| 2 | L | Superior Temporal Gyrus, posterior division | 2,008 | 4.40 | -64 | -12 | 2 |
| 3 | L | Frontal Medial Cortex | 1,104 | 4.75 | -8 | 42 | -12 |
| 3 | L | Frontal Pole | 1,104 | 4.38 | -6 | 58 | -16 |
| 3 | B | Frontal Medial Cortex | 1,104 | 4.05 | 0 | 42 | -16 |

**Table S14.** Descriptive statistics for clusters and local extrema showing a significant positive Valence × Temporal Certainty interaction during the acute presentation of faces and voices (Uncertain Threat—Uncertain Safety > Certain Threat—Certain Safety), *p*<.05, whole-brain corrected.

| Cluster | Hemisphere | Label | mm³ | *t* | *x* | *y* | *z* |
|---|---|---|---|---|---|---|---|
| 1 | L | Lateral Occipital Cortex, superior division | 5,160 | 5.18 | -14 | -66 | 60 |
| 1 | L | Precuneus Cortex | 5,160 | 5.26 | -8 | -64 | 52 |
| 1 | R | Precuneus Cortex | 5,160 | 5.36 | 10 | -58 | 56 |
| 2 | L | Pregenual anterior cingulate cortex | 4,408 | 4.53 | -2 | 38 | 12 |
| 2 | R | Pregenual anterior cingulate cortex | 4,408 | 4.81 | 4 | 36 | 14 |
| 2 | R | Frontal Medial Cortex | 4,408 | 4.57 | 10 | 50 | -8 |
| 2 | R | Frontal Pole/Dorsolateral Prefrontal Cortex | 4,408 | 4.38 | 8 | 58 | 8 |
| 2 | R | Paracingulate Gyrus | 4,408 | 3.54 | 6 | 34 | -10 |
| 3 | L | Postcentral Gyrus | 4,272 | 4.40 | -50 | -20 | 36 |
| 3 | L | Supramarginal Gyrus, anterior division | 4,272 | 5.21 | -62 | -30 | 32 |
| 4 | L | Precuneus Cortex | 1,912 | 4.75 | -16 | -68 | 24 |
| 5 | R | Angular Gyrus | 1,416 | 5.02 | 50 | -54 | 14 |
| 6 | L | Central Opercular Cortex | 1,336 | 3.58 | -44 | 6 | 0 |
| 6 | L | Insular Cortex | 1,336 | 4.80 | -38 | -8 | -10 |
| 7 | L | Frontal Medial Cortex | 984 | 4.73 | -10 | 48 | -10 |
| 7 | L | Frontal Pole/Dorsolateral Prefrontal Cortex | 984 | 3.96 | -4 | 58 | 0 |
| 8 | L | Inferior Frontal Gyrus, pars opercularis | 896 | 4.92 | -58 | 10 | 18 |
| 8 | L | Precentral Gyrus | 896 | 4.22 | -58 | 6 | 28 |
| 9 | R | Brain-Stem | 872 | 4.24 | 12 | -16 | -22 |
| 9 | R | Parahippocampal Gyrus, anterior division | 872 | 4.26 | 18 | -16 | -24 |
| 9 | R | Right Hippocampus | 872 | 4.43 | 26 | -22 | -18 |
| 9 | R | Temporal Fusiform Cortex, posterior division | 872 | 3.45 | 34 | -22 | -24 |

**Table S15.** Descriptive statistics for clusters and local extrema showing a significant negative Valence × Temporal Certainty interaction during the acute presentation of faces and voices (Uncertain Threat—Uncertain Safety < Certain Threat—Certain Safety), $p<.05$, whole-brain corrected.

| Cluster | Hemisphere | Label | mm³ | *t* | *x* | *y* | *z* |
|---|---|---|---|---|---|---|---|
| 1 | L | Intracalcarine Cortex | 65,456 | 12.80 | -16 | -72 | 6 |
| 1 | L | Lingual Gyrus | 65,456 | 8.43 | -18 | -68 | -4 |
| 1 | L | Occipital Pole | 65,456 | 7.24 | -6 | -98 | -4 |
| 1 | L | Cuneal Cortex | 65,456 | 6.62 | -2 | -88 | 34 |
| 2 | R | Intracalcarine Cortex | 65,456 | 12.58 | 18 | -68 | 8 |
| 1 | B | Lingual Gyrus | 65,456 | 11.50 | 0 | -88 | -4 |
| 1 | R | Supracalcarine Cortex | 65,456 | 11.24 | 2 | -74 | 12 |
| 1 | R | Cuneal Cortex | 65,456 | 7.38 | 8 | -84 | 36 |
| 1 | R | Occipital Pole | 65,456 | 7.09 | 12 | -104 | 2 |

**Table S16.** Descriptive statistics for clusters and local extrema showing greater activation during the uncertain versus certain presentation of threatening faces and voices (masked by the Valence × Certainty contrast), *p*<.05, whole-brain corrected.

| Cluster | Hemisphere | Label | $mm^3$ | *t* | *x* | *y* | *z* |
|---|---|---|---|---|---|---|---|
| 1 | L | Postcentral Gyrus | 1,640 | 4.04 | -50 | -20 | 34 |
| 1 | L | Supramarginal Gyrus, anterior division | 1,640 | 4.69 | -62 | -28 | 34 |
| 2 | R | Pregenual anterior cingulate cortex | 840 | 4.95 | 6 | 38 | 14 |
| 3 | L | Precentral Gyrus | 816 | 5.15 | -58 | 8 | 16 |
| 4 | L | Lateral Occipital Cortex, superior division | 696 | 3.62 | -14 | -68 | 56 |
| 5 | L | Precuneus Cortex | 672 | 4.74 | -16 | -66 | 24 |
| 6 | L | Central Opercular Cortex | 312 | 3.57 | -48 | 8 | 0 |
| 6 | L | Insular Cortex | 312 | 4.76 | -38 | -8 | -12 |

**Table S17.** Descriptive statistics for clusters and local extrema showing greater activation during the uncertain presentation of threatening versus benign faces and voices (masked by the Valence × Certainty contrast), $p$<.05, whole-brain corrected.

| **Cluster** | **Hemisphere** | **Label** | **mm³** | ***t*** | ***x*** | ***y*** | ***z*** |
|---|---|---|---|---|---|---|---|
| 1 | L | Pregenual anterior cingulate cortex | 1,752 | 5.04 | -2 | 36 | -4 |
| 1 | R | Pregenual anterior cingulate cortex | 1,752 | 4.66 | 6 | 40 | 10 |
| 2 | L | Central Opercular Cortex | 288 | 4.61 | -62 | -20 | 12 |
| 3 | R | Frontal Pole/Dorsolateral Prefrontal Cortex | 128 | 3.71 | 6 | 56 | 8 |
| 4 | L | Supramarginal Gyrus, anterior division | 120 | 3.31 | -64 | -38 | 24 |
| 5 | L | Paracingulate Gyrus | 40 | 3.64 | -8 | 46 | -2 |

**Table S18.** Descriptive statistics for clusters and local extrema showing greater activation during the uncertain presentation of threatening versus the certain presentation of benign faces and voices (masked by the Valence × Certainty contrast), $p$<.05, whole-brain corrected.

| **Cluster** | **Hemisphere** | **Label** | **mm³** | ***t*** | ***x*** | ***y*** | ***z*** |
|---|---|---|---|---|---|---|---|
| 1 | R | Brain-Stem | 8 | 3.60 | 10 | -18 | -20 |
| 2 | L | Precentral Gyrus | 8 | 3.28 | -58 | 6 | 32 |

**Table S19.** Descriptive statistics for clusters and local extrema showing greater activation during the uncertain presentation of benign versus the certain presentation of threatening faces and voices (masked by the Valence × Certainty contrast), $p$<.05, whole-brain corrected.

| **Cluster** | **Hemisphere** | **Label** | **mm³** | ***t*** | ***x*** | ***y*** | ***z*** |
|---|---|---|---|---|---|---|---|
| 1 | L | Precuneus Cortex | 968 | 4.97 | -12 | -64 | 26 |
| 2 | R | Precuneus Cortex | 712 | 4.33 | 8 | -58 | 52 |
| 3 | L | Precuneus Cortex | 8 | 3.32 | -2 | -62 | 18 |

**Table S20.** Descriptive statistics for clusters and local extrema showing greater activation during the certain presentation of benign versus the uncertain presentation of threatening faces and voices (masked by the Valence × Certainty contrast), *p*<.05, whole-brain corrected.

| **Cluster** | **Hemisphere** | **Label** | **mm$^3$** | ***t*** | ***x*** | ***y*** | ***z*** |
|---|---|---|---|---|---|---|---|
| 1 | L | Precuneus Cortex | 648 | 5.57 | -6 | -56 | 16 |
| 2 | L | Lateral Occipital Cortex, superior division | 408 | 4.06 | -16 | -66 | 64 |
| 3 | L | Frontal Medial Cortex | 64 | 4.51 | -6 | 46 | -12 |
| 4 | L | Precuneus Cortex | 8 | 3.30 | -8 | -66 | 22 |

**Table S21.** Descriptive statistics for clusters and local extrema showing greater activation during the certain presentation of benign versus threatening faces and voices (masked by the Valence × Certainty contrast), $p$<.05, whole-brain corrected.

| Cluster | Hemisphere | Label | mm³ | *t* | *x* | *y* | *z* |
|---|---|---|---|---|---|---|---|
| 1 | L | Lateral Occipital Cortex, superior division | 5,104 | 6.24 | -12 | -64 | 60 |
| 1 | L | Precuneus Cortex | 5,104 | 6.21 | -2 | -56 | 48 |
| 1 | R | Precuneus Cortex | 5,104 | 6.47 | 10 | -60 | 52 |
| 2 | L | Postcentral Gyrus | 2,440 | 4.31 | -58 | -24 | 40 |
| 2 | L | Supramarginal Gyrus, anterior division | 2,440 | 5.34 | -62 | -28 | 32 |
| 3 | L | Precuneus Cortex | 1,840 | 7.28 | -6 | -54 | 18 |
| 4 | R | Parahippocampal Gyrus, anterior division | 744 | 5.45 | 18 | -16 | -24 |
| 4 | R | Right Hippocampus | 744 | 6.18 | 26 | -20 | -18 |
| 5 | L | Frontal Medial Cortex | 472 | 4.19 | -8 | 48 | -10 |
| 5 | L | Frontal Pole/Dorsolateral Prefrontal Cortex | 472 | 3.61 | -4 | 56 | -2 |
| 5 | B | Frontal Medial Cortex | 472 | 3.69 | 0 | 52 | -8 |
| 6 | R | Frontal Medial Cortex | 336 | 4.66 | 10 | 54 | -8 |
| 7 | R | Angular Gyrus | 184 | 3.94 | 44 | -56 | 22 |
| 8 | R | Paracingulate Gyrus | 8 | 3.25 | 8 | 42 | -10 |

**Table S22.** Descriptive statistics for clusters and local extrema showing greater activation during the certain versus uncertain presentation of benign faces and voices (masked by the Valence × Certainty contrast), $p$<.05, whole-brain corrected.

| **Cluster** | **Hemisphere** | **Label** | **mm³** | ***t*** | ***x*** | ***y*** | ***z*** |
|---|---|---|---|---|---|---|---|
| 1 | L | Lateral Occipital Cortex, superior division | 1,248 | 4.76 | -14 | -66 | 62 |
| 1 | L | Precuneus Cortex | 1,248 | 3.77 | -8 | -58 | 62 |
| 2 | R | Pregenual anterior cingulate cortex | 1,040 | 4.02 | 6 | 40 | -2 |
| 2 | R | Frontal Medial Cortex | 1,040 | 5.05 | 10 | 50 | -8 |
| 3 | L | Frontal Medial Cortex | 984 | 5.88 | -6 | 48 | -12 |
| 4 | R | Angular Gyrus | 816 | 4.52 | 52 | -56 | 16 |
| 5 | L | Precuneus Cortex | 688 | 4.35 | -8 | -56 | 10 |
| 6 | R | Precuneus Cortex | 632 | 4.38 | 12 | -58 | 58 |
| 7 | L | Central Opercular Cortex | 232 | 3.97 | -62 | -18 | 14 |
| 8 | R | Frontal Pole/Dorsolateral Prefrontal Cortex | 152 | 4.14 | 10 | 62 | 0 |
| 9 | L | Pregenual anterior cingulate cortex | 104 | 4.13 | -4 | 36 | -6 |
